# MANF regulates unfolded protein response and neuronal survival through its ER-located receptor IRE1α

**DOI:** 10.1101/2020.09.22.307744

**Authors:** Vera Kovaleva, Li-Ying Yu, Larisa Ivanova, Jinhan Nam, Ave Eesmaa, Esa-Pekka Kumpula, Juha Huiskonen, Päivi Lindholm, Merja Voutilainen, Mati Karelson, Mart Saarma

## Abstract

Mesencephalic astrocyte-derived neurotrophic factor (MANF) is an endoplasmic reticulum (ER)-located protein with cytoprotective effects in numerous cell types *in vitro* and in models of neurodegeneration and diabetes *in vivo*. So far, the exact mode of its action has remained elusive and plasma membrane or ER-located receptors of MANF have not been identified. We have found that MANF can directly interact with transmembrane unfolded protein response (UPR) receptor IRE1α and compete with the major ER chaperone BiP (GRP78) for the interaction with IRE1α. With lower affinities MANF can also interact with other UPR receptors, PERK and ATF6. Using molecular modeling and mutagenesis analysis, we have identified the exact structural MANF regions involved in its binding to the luminal domain of IRE1α. MANF attenuates UPR signaling by decreasing IRE1α oligomerization and IRE1α phosphorylation. MANF mutant deficient in IRE1α binding cannot regulate IRE1α oligomerization and fails to protect neurons from ER stress induced death. Importantly, we found that MANF-IRE1α interaction is also crucial for the survival promoting action of MANF for dopamine neurons in an animal model of Parkinson’s disease. Our data reveal a novel mechanism of IRE1α regulation during ER stress and demonstrate the intracellular mode of action of MANF as a modulator of UPR and neuronal cell survival through the direct interaction with IRE1α and regulation of its activity. Furthermore, our data explain why MANF in contrast to other growth factors has no effects on naive cells and rescues only ER stressed or injured cells.

## Introduction

Endoplasmic reticulum (ER) is the largest intracellular compartment in most eukaryotic cells, dealing with protein secretion and folding as well as lipid biosynthesis and calcium homeostasis. Upon overloading of ER with misfolded proteins, a process occurring in many physiological and pathological conditions, a signaling machinery called unfolded protein response (UPR) is activated. The UPR aims for restoring cellular homeostasis through the activation of pro-survival signalling cascades, though when activated chronically it leads to apoptosis. UPR signalling in mammalian cells occurs through three ER transmembrane sensors: IRE1α (inositol-requiring enzyme 1α), PERK (PKR-like endoplasmic reticulum kinase) and ATF6 (activating transcription factor 6) (Walter and Ron, 2011). The activation of UPR sensors induces ER chaperones, triggers the protein degradation machinery and attenuates protein synthesis, thereby reducing the misfolded protein load in ER. IRE1 branch of UPR is the most evolutionarily conserved, present even in yeast cells (Cox et al., 1993). The major ER chaperone binding immunoglobulin protein (BiP), alias GRP78, is classically believed to prevent IRE1α activation and signaling in basal conditions. The dissociation of BiP from luminal domain of IRE1α upon ER stress leads to dimerization/oligomerization of luminal domains of IRE1α, resulting in trans-autophosphorylation of cytoplasmic domains of IRE1α, increasing IRE1α endoribonuclease activity and triggering unconventional splicing of X-box-binding protein 1 (XBP1) mRNA (Bertolotti et al., 2000). IRE1α is a unique transmembrane receptor where the cytoplasmic domain is an enzyme, possessing both serine-threonine kinase and endoribonuclease activities. As endoribonuclease, it cuts out intron from mRNA of X-box-binding protein 1 (XBP1), triggering UPR activation. Through its endoribonuclease activity, IRE1α also causes mRNA decay, allowing to decrease the protein synthesis load during ER stress. Phosphorylation of IRE1α within the kinase activation loop results in increased endoribonuclease activity (Prischi et al., 2014). Despite the fact that IRE1 being discovered almost 30 years ago, the exact mechanism of its activation is not entirely clear. There are a few mutually exclusive theories, supporting different modes of IRE1α activation (Karagöz et al., 2017; Amin-Wetzel et al., 2017; Preissler and Ron, 2018; Carrara et al., 2015b; Kopp et al., 2018). Recently unfolded proteins and chaperones such as Heat shock protein 47 (Hsp47) and protein disulfide isomerase A6 (PDIA6) were shown to play a role in the regulation of IRE1α activation (Sepulveda et.al, 2018, Eletto et al., 2014). Unfolded proteins were shown to bind IRE1α LD, and through the induction of allosteric changes promote IRE1α oligomerization (Karagöz et al., 2017). Heat shock protein 47 (Hsp47) enhances IRE1α activation through direct interaction and displacement of IRE1α attenuator BiP from the complex (Sepulveda et.al, 2018). Protein disulfide isomerase A6 (PDIA6) has been shown to attenuate IRE1α signaling upon ER stress after BiP dissociation from IRE1α through the reduction of disulfide bonds formed by Cys148 residues from individual IRE1α monomers in the oligomeric IRE1α, thus converting IRE1α back to monomeric form (Eletto et al., 2014).

Mesencephalic astrocyte-derived neurotrophic factor (MANF) (Petrova et al., 2003) together with cerebral dopamine neurotrophic factor (CDNF) (Lindholm et al., 2007) forms a novel family of evolutionary conserved ER-located, but also secreted unconventional neurotrophic factors (Lindahl et al., 2017). In animal models MANF promotes the survival of dopamine neurons, which degenerate in Parkinson’s disease (PD) (Voutilainen et al., 2009) and protects pancreatic beta cells from death (Lindahl et al., 2014, Danilova et al., 2019). MANF is up-regulated in ER stress conditions, bypassing general down-regulation of protein synthesis (Apostolou et al, 2008). MANF has been shown to protect ER stressed cells and alleviate UPR markers in a number of *in vitro* models (Mizobuchi et al., 2007; Apostolou et al., 2008; Tadimalla et al., 2008; Hellmann et al., 2011; Pakarinen et al., 2020). Recent data also show that MANF is a key regulator of metabolic and immune homeostasis in ageing. Moreover, MANF protects against liver inflammation and fibrosis, suggesting a therapeutic application for MANF in age-related metabolic diseases (Sousa-Victor et al. 2019). In MANF-deficient mice UPR pathways are chronically activated in beta cells, in neurons and several other cell types demonstrating that MANF is a crucial regulator of UPR *in vivo* (Lindahl et al., 2014; Danilova et al., 2019; Pakarinen et al., 2020).

To date, the exact mode of action of this protein remains poorly understood. According to our previously published data, in human beta-cells MANF is mostly localized inside the cells in the ER (Danilova et al., 2019), which implies that ER is its main site of action. In line with this MANF added extracellularly to superior cervical ganglion (SCG) neurons had no prosurvival effect, but when the plasmid encoding MANF or MANF protein were microinjected they protected the neurons from apoptosis, including ER stress induced apoptosis (Hellmann et al., 2011; Mätlik et al., 2015; Eesmaa et al., under review). It is not clear how MANF exerts its protective effects mostly acting in the ER lumen. Since MANF can be secreted it acts extracellularly as well but this mechanism is also entirely unclear. In ER stress MANF secretion increases and MANF added extracellularly to cultured pancreatic beta cells has clear cytoprotective and proliferative effect (Lindahl et al., 2014), showing that MANF can act via unknown plasma membrane receptors or enter the cells, translocate to the ER and act there. MANF interacts with major ER chaperone BiP (Glembotski et al, 2012) and other chaperones, including PDIA6 (Bell at al., 2019; Eesmaa et al., under review). Recently MANF was shown to prolong the interaction of BiP with client proteins, thus regulating protein-folding homeostasis (Yan et al. 2019). All these findings support the hypothesis that the major locus operandi of MANF is the ER lumen.

Interestingly, MANF knockout mice similarly to both IRE1α or XBP1 knockout mice have diabetic-like phenotypes. All three knockout models have endocrine pancreas alterations, altered glucose metabolism and insulin secretion, as well as lipid abnormalities in liver (Bommiasamy & Popko, 2011; Hetz et al., 2012; Lindahl et al., 2014; Danilova et al., 2019; Sousa-Victor et al. 2019). These data provide genetic evidence for MANF and IRE1α being involved in similar functions and signaling pathways in cells. MANF-IRE1α crosstalk is further supported by the notion that in MANF knockout mice IRE1α branch of UPR is activated first, and PERK and ATF6 pathways are activated later (Lindahl et al., 2014). Upregulation of spliced XBP1 (sXBP1) downstream to IRE1α was shown to occur first not only for full knockout mice but also for pancreas-specific (Danilova et al., 2019) and for central nervous system-specific MANF ablation (Pakarinen et al., 2020).

We have recently shown that in ER stressed mouse cultured dopamine neurons MANF is able to reduce the expression of *sXbp1, Atf6* as well as *Bip* mRNA. In a similar way, chemical inhibition of both IRE1α and PERK pathways abolishes the anti-apoptotic effect of MANF in mouse sympathetic neurons and in dopamine neurons (Eesmaa et al., under review). These data indicate that UPR pathways are involved in the pro-survival action of MANF. Interestingly, the interaction with BiP was not required for the prosurvival activity of MANF (Eesmaa et al., under review). Considering the protective effects of MANF in ER stressed cells, its interaction with several chaperones and the genetic evidence, we tested the hypothesis that MANF directly binds to IRE1α and through that regulates the IRE1α branch of UPR.

Here we show that MANF directly binds to IRE1α with high affinity and also interacts with PERK and ATF6 with lower affinities. We found that MANF competes with BiP for the interaction with IRE1α. We confirm that through the direct interaction MANF regulates UPR by inhibiting the activity of IRE1α through regulating its phosphorylation, oligomerization and downstream signaling. MANF mutant deficient in IRE1α binding is unable to regulate IRE1α activity and lacks pro-survival action *in vitro* in SCG and dopamine neurons and is not biologically active *in vivo* in an animal model of Parkinson’s disease. Thus, our results reveal the mode of action of MANF in the ER and bring to light MANF as a novel regulator of IRE1α activity.

## Results

### MANF directly interacts with luminal domain of IRE1α

To investigate the mechanism how MANF is regulating UPR and crosstalking with the UPR machinery, we started by testing for direct interaction between MANF and luminal domains (LDs) of UPR sensors. We expressed and purified from *CHO* cells the luminal domains of all three UPR sensors, human IRE1α, PERK and ATF6. We confirmed that the proteins are highly pure and intact (Supplementary Fig. 1a). We also tested their glycosylation status using peptide N-Glycosidase F (PNGase F) assay. We found that IRE1α LD and PERK LD were not N-glycosylated, whereas we confirmed the earlier findings that ATF6 LD was glycosylated (Supplementary Fig. 1a). Glycosylated glial cell line-derived neurotrophic factor (GDNF) served as the positive control for the assay. To assess the biological activity of LD of UPR sensors we tested their interactions with their known binding partner BiP using microscale thermophoresis (MST) and purified recombinant proteins. We found that BiP was interacting with fluorescently labeled through His-tag UPR sensors with high affinities: BiP-IRE1α LD K_d_=119.3±34.0 nM, BiP-PERK LD K_d_=14.1±11.5 nM and BiP-ATF6 LD K_d_=21.0±15.2 nM (Fig. 1a, b, c). Notably, the affinities of BiP for these mammalian cell produced UPR receptors were about 10-100 times higher than those reported for *E*.*coli* produced proteins (Cararra et. al., 2015).

**Fig. 1.**
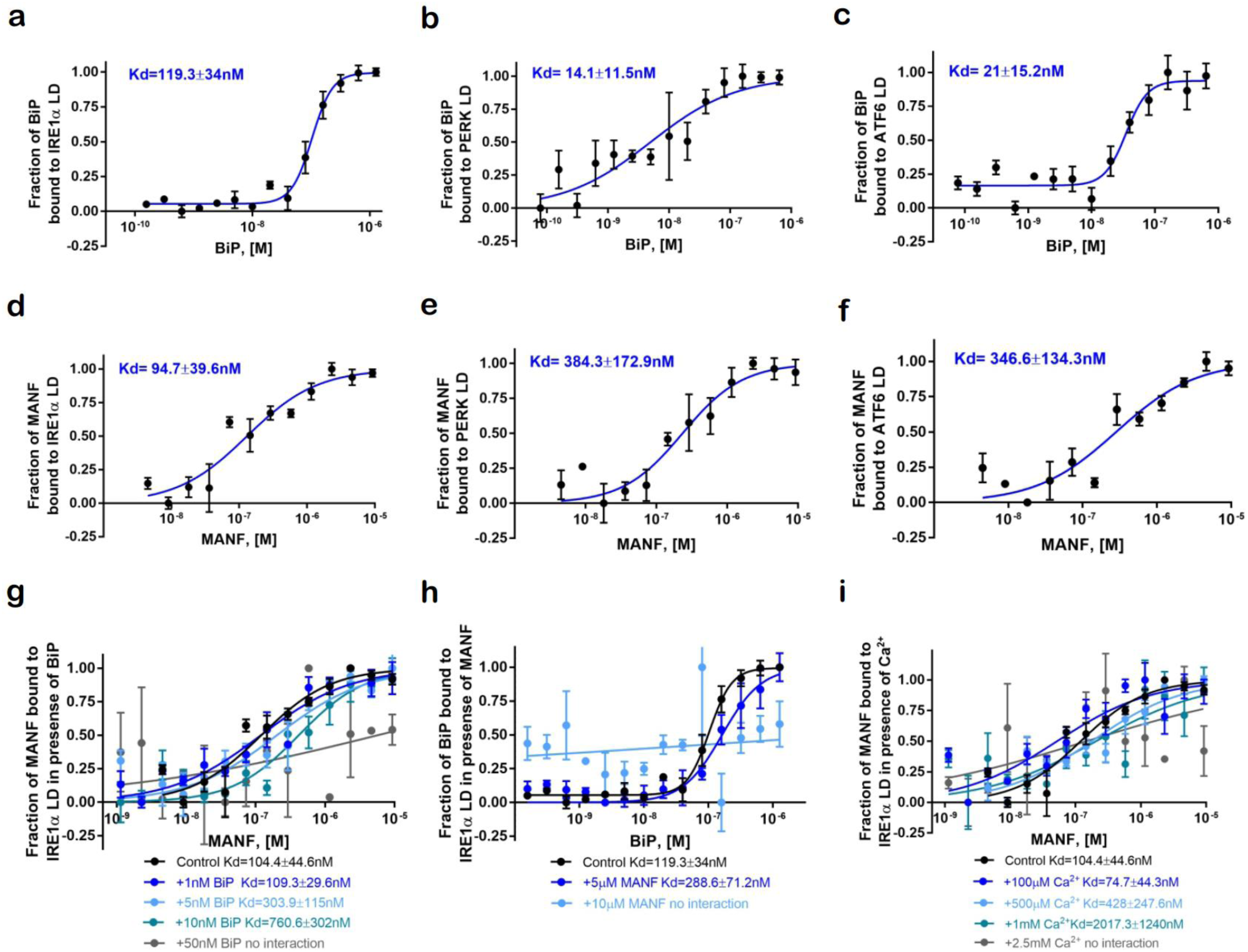
MANF directly interacts with luminal domain of IRE1α and BiP is preventing MANF interaction with IRE1α. **a, b, c**, Labeled through His-tag luminal domains (LDs) of IRE1α, PERK, ATF6 (20nM) interact with unlabeled titrated recombinant purified BiP protein (0-640 nM), as shown using microscale thermophoresis (MST). **d, e, f**, Purified recombinant MANF protein (0-9.3 µM) is interacting with labeled through His-tag LDs of UPR sensors IRE1α, PERK, ATF6 (20nM). **g**, MANF-IRE1α LD interaction in presence of BiP(1-50 nM). Purified recombinant MANF protein is titrated (0-9.3 µM), and IRE1 LD concentration is 20nM. **h**, BiP-IRE1α LD interaction is in presence of 5µM (10 µM) of purified MANF protein (10 nM-1µM). Purified BiP protein is titrated (0-640 nM) and incubated with Alexa647-labeled through His-tag luminal domain of IRE1α (20nM). **i**, Interaction of unlabeled titrated human recombinant MANF (0-9.3 µM) with Alexa647-labeled through His-tag luminal domain of IRE1α (20nM) in presence of increasing concentrations of Ca^2+^ concentration (100µM-2.5mM). Microscale thermophoresis binding curves, showing mean fraction bond values from n=3-5 experiments per binding pair ±SEM, Kd values±error estimations are indicated.

To determine whether there is a direct interaction between MANF and LDs of UPR sensors, we used mammalian cell-line produced recombinant purified human MANF and LDs of IRE1α, PERK and ATF6 (Supplementary Fig. 1a). MANF prepared this way was biologically active, as shown by survival assay in SCG sympathetic neurons and dopamine neurons (Eesmaa et al., under review). Fluorescently labeled through His-tag LDs of UPR sensors were incubated with increasing concentrations of MANF and the binding affinities were measured using MST. We observed that MANF is interacting with all three UPR sensors, and the highest affinity K_d_=94.7±39.6 nM was observed for MANF-IRE1α LD interaction (Fig. 1d). MANF was interacting with PERK and ATF6 LDs with lower affinities, K_d_=384.3±172.9 nM and, K_d_=346.6±134.3 nM, respectively (Fig. 1e, f). This is the first demonstration of direct interactions between MANF and UPR sensors.

Taking into account the similarities in MANF and IRE1α knockout mouse phenotypes, early activation of IRE1α branch of UPR in MANF knockout mice and a high affinity for MANF to IRE1α LD, as compared to PERK and ATF6 LDs, we decided to further focus on MANF-IRE1α LD interaction. We tested the interaction between purified recombinant *CHO* produced MANF and IRE1α LD using two other methods in addition to MST. In gel filtration chromatography the MANF-IRE1α LD complex was co-purified and revealed that MANF interacts with the monomer of IRE1α LD (Supplementary Fig. 1b). In binding on nickel coated plates followed by ELISA-based MANF detection we again confirmed that MANF binds to IRE1α LD-His (Supplementary Fig. 1c).

### BiP prevents MANF interaction with IRE1α LD

As shown above, both MANF and BiP were able to bind IRE1α LD with similar affinities. Therefore, we hypothesised that BiP can prevent the interaction between MANF and IRE1α LD if MANF and BiP compete for the same binding site on IRE1α LD. Alternatively, MANF binding to BiP may increase the affinity of BiP interaction with IRE1α allosterically, if they form a tripartite complex. To distinguish between these two hypotheses we tested the interaction between MANF and IRE1α LD in presence of increasing concentrations of BiP (1nM-50nM) using MST. We found that 10nM BiP decreased the affinity of MANF binding to IRE1α LD and 50nM BiP abolished the interaction (Fig. 1g). This result suggests that MANF and BiP compete for the same binding site on IRE1α LD. To investigate whether BiP abolished the MANF-IRE1α LD interaction due to MANF-BiP interaction, we tested purified recombinant human MANF mutant proteins MANF E153A and MANF R133E deficient in BiP binding (Yan et al., 2019; Eesmaa et al., under review) for binding to IRE1α LD. We found that MANF E153A and MANF R133E were binding IRE1α LD with similar affinities as wild-type MANF, confirming that binding sites for BiP and IRE1α LD are different (Supplementary Fig. 1d). In addition, we also tested whether BiP affects the binding of MANF E153A and MANF R133E to IRE1α LD and found that similarly to wild-type MANF 50nM BiP abolishes the interaction of BiP binding deficient MANF mutants with IRE1α LD (Supplementary Fig. 1e). These results demonstrate that, BiP-MANF interaction does not affect BiP competition with MANF for IRE1α LD binding. We conclude that BiP either competes with MANF for the same binding interfaces on the LD of IRE1α or changes the conformation of the LD to being restrictive for MANF binding.

We then tested whether MANF can increase the affinity of BiP to IRE1α LD. Interestingly, in the presence of increasing concentrations of MANF (10 nM-1 µM) BiP was still able to interact with IRE1α LD (Supplementary Fig. 1f). At unphysiological 10 µM concentrations of MANF the BiP-IRE1α LD interaction was abolished (Fig. 1h). Based on these data we hypothesised that MANF can bind IRE1α LD only when BiP is not bound to IRE1α, as it happens in conditions of ER stress, when BiP dissociates from UPR sensors to bind misfolded or aggregated proteins. We favor the interpretation that BiP binding changes the conformation of IRE1α LD so that it loses the high affinity binding site for MANF, because MANF is not displacing BiP from the complex with IRE1α at the concentration equal to its K_d_ to IRE1α.

### Ca^2+^ regulates MANF-IRE1α LD interaction

Considering that ER is crucial for the maintenance of cellular calcium homeostasis and Ca^2+^ depletion from ER is known to activate UPR, we have tested whether Ca^2+^ affects MANF interaction with UPR sensors. The concentration of free Ca^2+^ in ER has been reported to be between low µM range to up to 1-3 mM, depending on the techniques used for measurements as well as cell types (Zampese & Pizzo, 2012). We started from testing of interactions between MANF and UPR sensors in the presence of 500 µM Ca^2+^, since this concentration is most often reported as a physiologically relevant concentration of free Ca^2+^ in ER in basal conditions. At 500 µM Ca^2+^ the affinity of MANF to IRE1α LD was slightly decreased and no changes in the affinities of MANF to PERK and ATF6 LDs were found (Supplementary Fig. 1g, h, i). When MANF-IRE1α LD interaction was further tested in presence of increasing concentrations of Ca^2+^ (100 nM-2.5 mM), we found that the affinity of MANF to IRE1α was inversely proportional to Ca^2+^ concentration (Fig. 1i). Since Ca^2+^ is depleted from the ER lumen upon ER stress (Mekahli et al., 2011), decrease in Ca^2+^ concentration in the ER can facilitate MANF-IRE1α interaction in ER-stressed, but not in naive cells.

### C-terminal domain of MANF directly interacts with IRE1α LD

Since we have earlier found that 63 amino acid C-terminal domain of MANF (C-MANF) can protect neurons from chemically induced cell death as efficiently as full-length MANF (Hellman et al, 2011), we hypothesized that binding of MANF to IRE1α may occur through C-MANF. We tested the interactions between chemically synthesized or *E*.*coli* produced C-MANF with luminal domains of human IRE1α, PERK and ATF6 as it was done for full-length MANF. We showed that chemically synthesized C-MANF is interacting with all three UPR sensors with higher affinities as compared with full-length MANF: C-MANF-IRE1α LD K_d_=9.5±4.2 nM, C-MANF-PERK LD K_d_=7.6±3.1 nM, C-MANF-ATF6 K_d_=9.8±4.3 nM (Fig. 2a, b, c). We also tested chemically synthesized C-MANF in a survival assay in SCG sympathetic neurons and found it biologically active, i.e. possessing the ability to promote survival SCG neurons in the absence of nerve growth factor (NGF) (Fig. 2d). We showed that chemically synthesized C-MANF is homogeneous using high-performance liquid chromatography (HPLC) and has the required disulfide bond between Cys128 and Cys130 using mass spectrometry (MS) (Supplementary Fig. 2a, b). We found that *E*.*coli* produced C-MANF is also interacting with all three UPR sensors, but the affinities were significantly lower as compared with full-length MANF and chemically synthesized C-MANF. As with full-length MANF, *E*.*coli* produced C-MANF also had the highest affinity to IRE1α LD K_d_=241.2±107.5 nM and lower affinities to PERK and ATF6 LDs, K_d_=1.4±0.6 µM and K_d_=1.3±0.7 µM, correspondingly (Supplementary Fig. 2c). These data confirm our finding that *E*.*coli* synthesized MANF and its variants generally have lower affinity for the LDs of UPR sensors and perhaps also other binding partners.

**Fig. 2.**
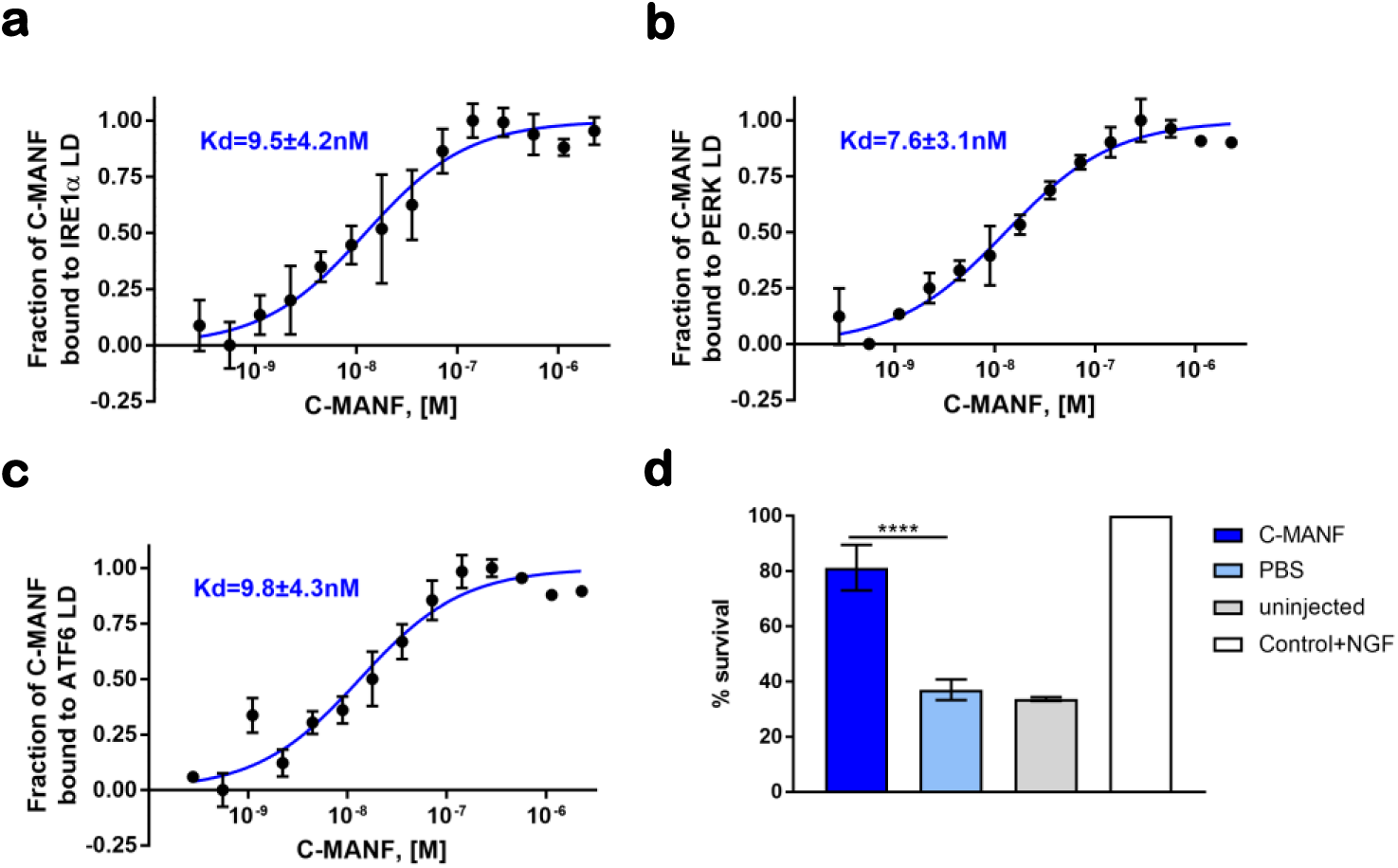
C-MANF directly interacts with luminal domain of IRE1α. **a, b, c**, Chemically synthesized C-MANF (0-2.28 µM) is interacting with labeled through His-tag LDs of UPR sensors IRE1α, PERK, ATF6 (20nM). Microscale thermophoresis binding curves, showing mean fraction bond values from n=3-4 experiments per binding pair ±SEM, Kd values±error estimations are indicated. **d**, Microinjections of C-MANF protect sympathetic SCG neurons upon nerve growth factor (NGF) deprivation, n=3.

### MANF interacts with the luminal domain of IRE1α in cells

To test whether the interaction between MANF and IRE1α also occurs in cells, we performed *in situ* proximity ligation assay (PLA) in Flp-In-TREx293 and CHO cells. In the first set-up, we investigated the interaction between endogenous MANF and overexpressed IRE1α with hemagglutinin tag (IRE1α-HA) in isogenic stable cell lines, expressing BiP-HA (positive control), IRE1α-HA, GFP-HA (negative control) upon doxycycline induction. We found that in Flp-In-TREx293 cells endogenous MANF interacts with IRE1α-HA (Fig 3a, b). In addition, we tested MANF-IRE1α interaction in CHO cells, in order to analyse the interaction between endogenous IRE1α and overexpressed MANF-HA We found that endogenous IRE1α interacts with MANF-HA in CHO cells (Supplementary Fig. 3a)., MANF-HA construct was biologically active in a survival assay in SCG neurons (Eesmaa et al., under review). As an additional to GFP-HA negative control we used a cell line expressing truncated version of MANF, lacking a major part of the C-terminal domain (Supplementary Fig. 3b).

**Fig. 3.**
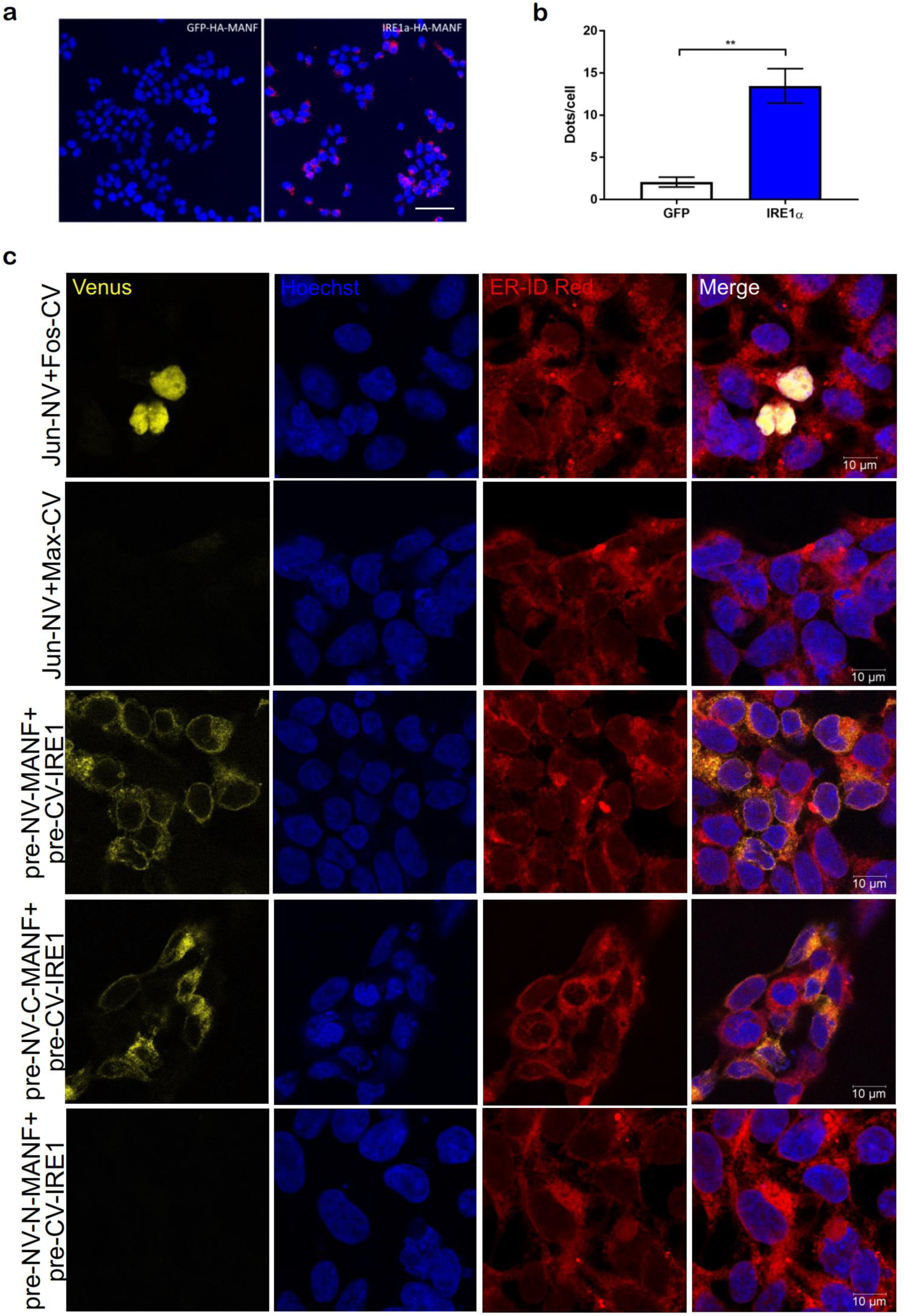
MANF interacts with IRE1α in cells. **a**, Representative image of MANF-IRE1α-HA interaction in Flp-in TREx293 cells, Flp-in TREx293 GFP-HA was used as a negative control. **b**, Quantification of dots per cell for MANF-IRE1α-HA interaction versus MANF-GFP-HA interaction. Mean dots/cell values ±SEM from n=3 independent experiments are indicated. Statistical analysis was performed using Student’s t test: **: p < 0.01. **c**, MANF interacts with IRE1α through its C-terminal domain but not N-terminal domain in HEK293 cells, as shown using bimolecular fluorescence complementation assay (BiFC), n=3. Statistical analysis was performed using one-way ANOVA, followed by Holm-Sidak’s multiple comparison test, ****: p < 0.0001.

We further verified the interaction between MANF and IRE1α LD in cells using bimolecular fluorescence complementation assay (BiFC). HEK293 cells were transfected with bait and prey proteins fused either to the N-terminal (NV) or C-terminal fragment of Venus, a variant of yellow fluorescent protein (YFP). If bait and prey proteins are interacting, the split Venus fragments reconstitute and form full-length fluorescent protein, enabling the detection of interaction using fluorescence microscopy. To ensure the correct cellular localization of tested proteins we designed the constructs with Venus fragments located between the signal sequences and mature proteins. As positive controls we used transcription factors Jun and Fos and known interactor of IRE1α LD BiP. To check for the absence of non-specific background signal, we tested also the interaction between Jun and Max, two non-interacting transcription factors.

Using BiFC, we demonstrated that pre-NV-MANF is interacting with pre-CV-IRE1α LD in HEK293 cells (Fig. 3c). Interestingly, we found that the interaction between MANF and IRE1α LD in cells occurs through C-terminal domain of MANF (C-MANF). Moreover, N-terminal domain of MANF (N-MANF) was not interacting with IRE1α LD (Fig. 3c). Since it is currently debatable which of the domains of BiP is interacting with IRE1α LD, we tested the interaction of ATP-binding domain of BiP (BiP NBD) and substrate-binding domain of BiP (BiP SBD) with IRE1α LD. According to our data BiP mainly interacts with IRE1α LD through its NBD and for BiP SBD much weaker and less frequent signal was detected using BiFC (Supplementary Fig. 3c).

### Dissection of IRE1α LD binding site in MANF

We confirmed the interaction between MANF and IRE1α LD both for purified recombinant proteins and in cells using several techniques. We then set out to identify (predict) the exact binding sites between these proteins using known structures of MANF (PDB ID: 2KVD)/ C-MANF (PDB ID: 2KVE) and IRE1α LD (PDB ID: 2HZ6) (Fig. 4a) and computational modelling, followed by MANF site-directed mutagenesis and testing of the IRE1α LD binding and biological activity of the respective MANF mutants.

**Fig. 4.**
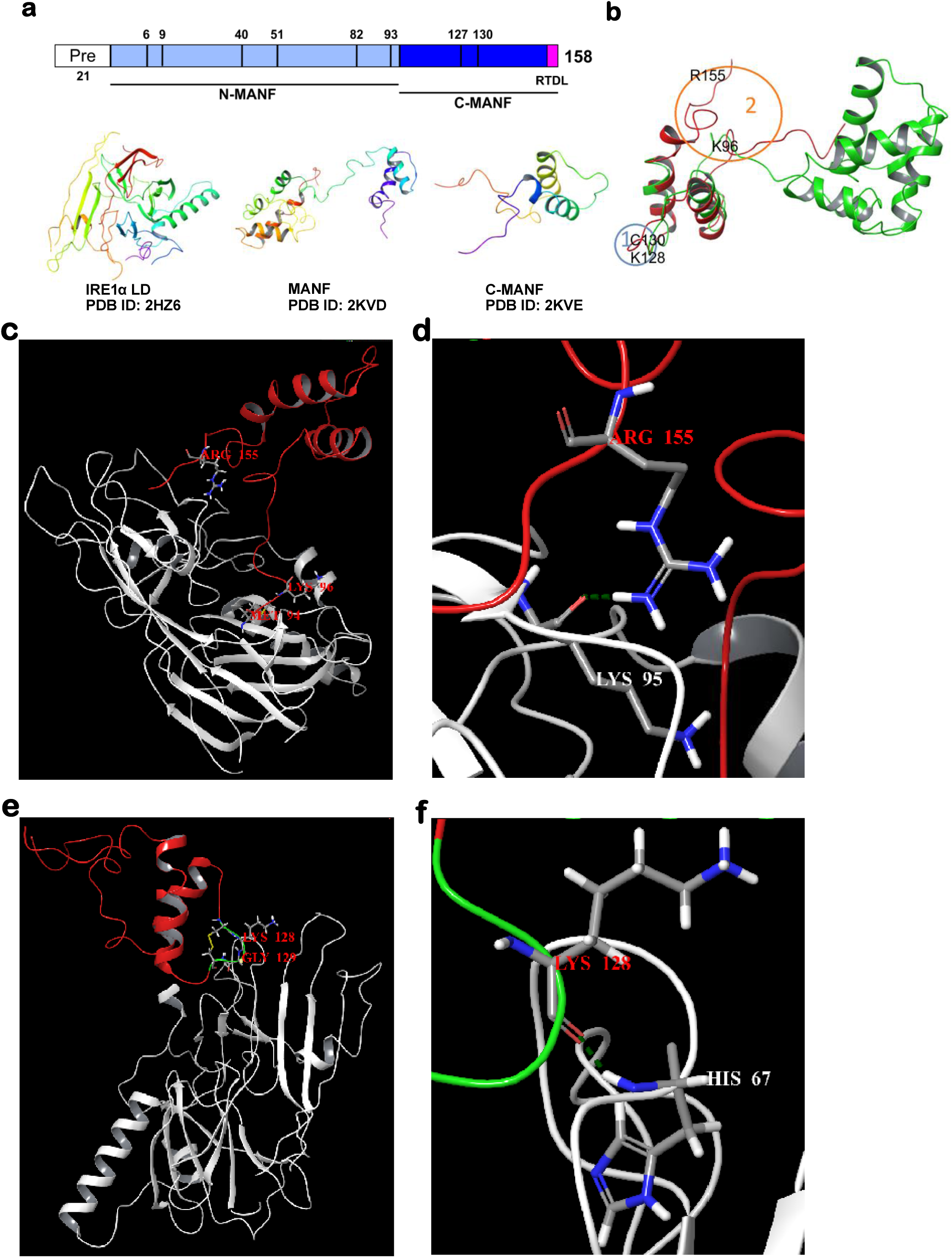
Putative MANF-IRE1α binding sites, predicted using molecular dynamics and molecular docking. **a**, Scheme of two-domain MANF structure and PDB structures used for computational modeling. **b**, Putative IRE1α binding regions 1 and 2 in the aligned structures of the C-terminal of MANF (pdb:2KWE, red ribbon) and whole MANF (pdb:2W51, green ribbon). **c**, Relative position of MANF and IRE1α proteins in complex configuration 12. **d**, The hydrogen bond between Arg155 amino acid residue of MANF and Lys95 amino acid residue of IRE1α proteins in complex configuration 12 (hydrogen bond is represented as green dashed line). **e**, Relative position of MANF and IRE1α proteins in complex configuration 41. The cysteine loop of the MANF is given in green color. **f**, The hydrogen bond between Lys128 amino acid residues of MANF and His67 amino acid residue of IRE1α proteins in complex configuration 41 (hydrogen bond is represented as green dashed line).

At first, molecular docking calculations were carried out for C-MANF in complex with IRE1α LD without any mutual protein position constrains. The IRE1α LD was considered acting as a receptor and MANF as a ligand and was docked flexibly. In result, 30 computationally generated protein-protein complex configurations (hereinafter referred as configurations) referring to different relative positions of MANF and IRE1α LD were obtained. The analysis of these configurations showed that IRE1α LD is contacted primarily by the link between C- and N-terminal domains of MANF protein involving the M94 - T105 and by K150 - L158 regions (Supplementary Table 1, Complex configurations 1-30).

**Table 1.** The analysis of the frequency of the appearance of each interacting amino acid residue in computational complex configurations 1-30.

| MANF amino acids residues | Interactions |  |  |
| --- | --- | --- | --- |
|  | Total | Total % | Hydrogen bonds |
| Met94 | 26 | 86,7 | 10 |
| Gly95 | 11 | 36,7 | 0 |
| Lys96 | 9 | 30,0 | 3 |
| Tyr97 | 27 | 90,0 | 2 |
| Asp98 | 24 | 80,0 | 3 |
| Lys99 | 29 | 96,7 | 0 |
| Gln100 | 26 | 86,7 | 4 |
| Ile101 | 24 | 80,0 | 0 |
| Ser104 | 17 | 56,7 | 2 |
| Thr105 | 16 | 53,3 | 2 |
| Lys150 | 22 | 73,3 | 1 |
| Ser153 | 10 | 33,3 | 1 |
| Ala154 | 14 | 46,7 | 3 |
| Arg155 | 23 | 76,7 | 7 |
| Thr156 | 19 | 63,3 | 3 |
| Asp157 | 18 | 60,0 | 2 |
| Leu158 | 23 | 76,7 | 0 |

In order to estimate the importance of individual amino acid residues of MANF in the interaction with IRE1α LD, the frequency of the appearance of each amino acid residue in 30 docking computational configurations was determined. Furthermore, the number of configurations involving hydrogen bonding at the complex interface was assessed for these individual MANF amino acid residues (Table 1).

The examination of the computational results led to the identification of three amino acids having the most frequent involvement in specific interactions, i.e. hydrogen bonds, the residues M94, K96 and R155 (Fig. 4b, c, d). The mutation at M94 is expected to have minor effect as it involves the hydrogen bond of the protein backbone. Nevertheless, all these three amino acids were proposed as targets for mutations.

The second series of molecular docking calculations was carried out with structural constraints at the MANF protein. Notably, MANF is a two-domain bifunctional protein (Parkash et al., 2009; Hellman et al., 2011) (Fig. 4a) and in principle its activity as a ligand for IRE1α can be supported by interactions involving amino acid residues from either N-terminal or C-terminal domain, or both. The disulfide bridge containing cysteine loop (C127-K128-G129-C130) in the C-terminal domain of MANF has been suggested as the most attractive site for binding, since earlier in (Mätlik et al., 2015) it was shown to be crucial for the anti-apoptotic activity of MANF in sympathetic SCG neurons. The analysis of the 30 configurations of MANF with IRE1α LD with the requirement of the involvement of the cysteine loop in the protein-protein binding was accordingly carried out as described above. Notably, only four configurations out of 30 featured significant interactions between the proteins. In this analysis, the frequency of appearance as binding residue in the configurations and the frequency of appearance in hydrogen bonds were hence calculated only for C127, K128, G129 and C130 amino acid residues of MANF (Table 2). The interactions between MANF and IRE1α LD in all configurations regarding the required involvement of the cysteine loop are listed in Table S2 in the supplementary materials (Complex configurations 31-60). Because of the suggested importance in the protein-protein interactions, all four amino acids in the disulfide bridge containing loop were still included as targets for mutations (Fig. 4b, e, f).

**Table 2.** Analysis of the frequency of the appearance of interacting amino acid residues of the disulfide bridge of the MANF in computational complex configurations 31-60.

| MANF amino acids residues | Interactions |  |  |
| --- | --- | --- | --- |
|  | Total | Total % | Hydrogen bonds |
| Cys127 | 7 | 23,3 | 0 |
| Lys128 | 7 | 23,3 | 2 |
| Gly129 | 6 | 20,0 | 1 |
| Cys130 | 7 | 23,3 | 0 |

In conclusion, the analysis of all computational results led to the identification of four potential binding regions in IRE1α LD (Supplementary Fig. 4a). In the case of MANF, only one additional potential binding site apart from the cysteine loop was identified (Fig. 4b, Supplementary Fig. 4b). Notably, proposed IRE1α LD binding regions in MANF are highly evolutionary conserved (Supplementary Fig. 4c).

### Expression of MANF mutants putatively deficient for IRE1α binding

Following the structural predictions of MANF-IRE1α LD interactions from the computational modelling we generated MANF mutants, putatively deficient in IRE1α LD binding. The cysteine and lysine/cysteine in cysteine loop were mutated to serine and alanine/serine, correspondently (C130S and K128A C130S), the lysine in linker region between N- and C-terminal domains and arginine in RTDL-sequence were mutated to alanines (K96A and R155A). The levels of expression and secretion of generated mutant constructs were determined in CHO cells and were similar to that of the wild type MANF construct (wtMANF) (Supplementary Fig. 5a).

We also investigated the cellular localization of MANF mutants using immunocytochemistry and microinjections of plasmids encoding MANF mutants in mouse SCG neurons. No changes in the localization of MANF mutants compared to wtMANF were observed (Supplementary Fig. 5b, c).

### MANF reduces IRE1α oligomerization upon ER stress

Upon the loading of ER with misfolded proteins dissociation of BiP from luminal domain of IRE1α is thought to be the trigger of IRE1α dimerization, phosphorylation and activation (Preissler and Ron, 2018). Also binding of unfolded proteins can trigger IRE1α oligomerization and activation (Karagöz et al., 2017).

Since luminal domain of IRE1α interacting with BiP and misfolded proteins clearly determines its oligomeric status and activation, and MANF binds to IRE1α LD with high affinity, we decided to assess the effect of MANF on the oligomerization of IRE1α. For this, we have generated and used a doxycycline inducible stable Flp-In TREx-293 cell line expressing GFP-tagged IRE1α (Li et. al., 2010) (Supplementary Fig. 6a). Upon the treatment with ER stressor, we can monitor and quantify the level of IRE1α oligomerization, by measuring the number of IRE1α-GFP foci per cell. We found that transient 48-hour overexpression of MANF decreased the number of IRE1α-GFP foci per cell upon the treatment with the inhibitor of N-linked glycosylation and protein folding ER stressor tunicamycin (TM) (Fig. 5a). Expression of empty vector had no effect on IRE1α oligomerization and served as a negative control.

**Fig. 5.**
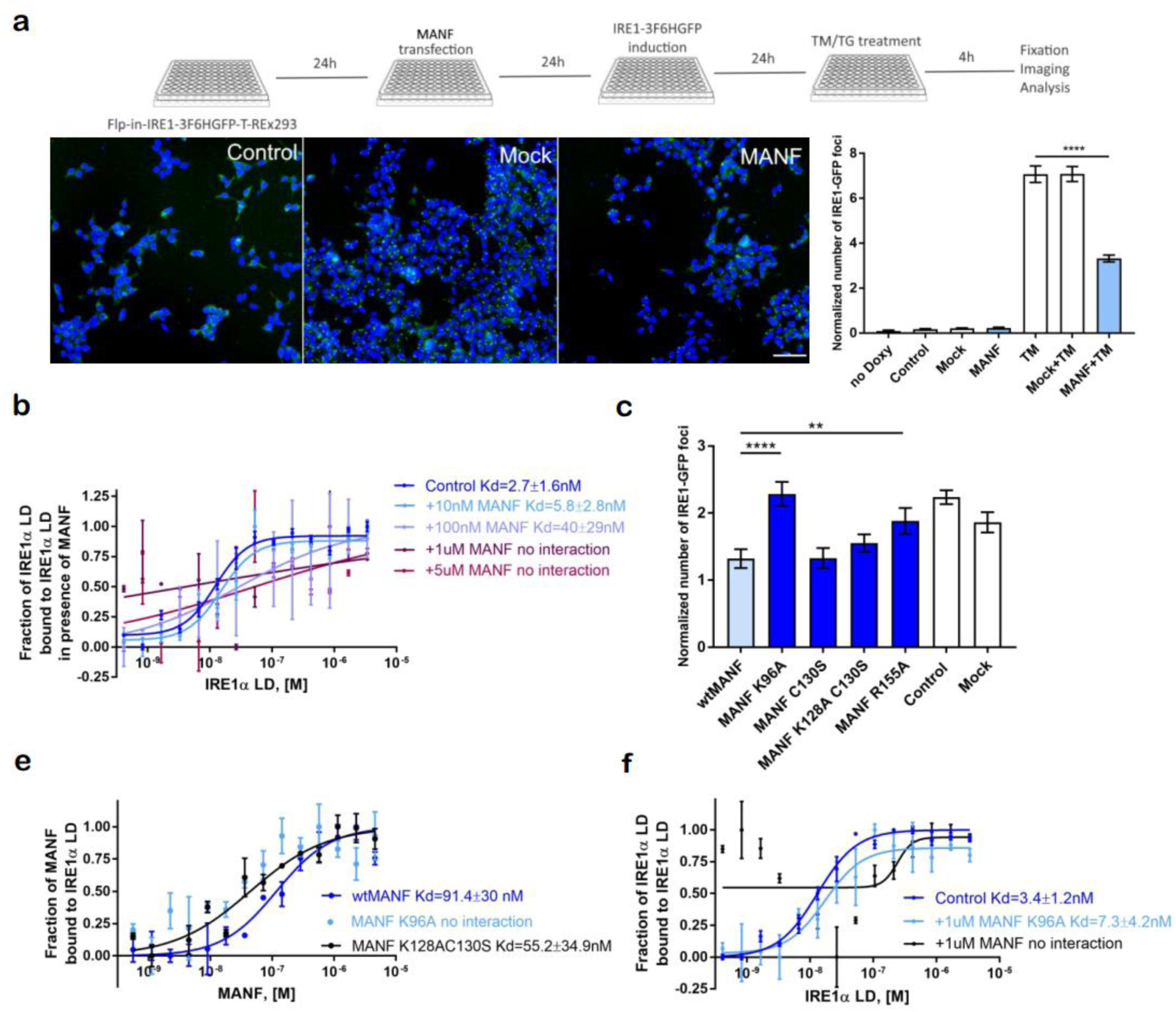
MANF is decreasing IRE1α oligomerization and IRE1α binding deficient MANF mutant is not affecting IRE1 α oligomerization upon ER stress. **a**, MANF-overexpression is decreasing IRE1α oligomerization upon ER stress, timeline of experiment and representative image. A stable Flp-In293 T-REx cell line expressing reporter IRE1α-3F6HGFP upon doxycycline induction was transfected with hMANF cDNA expressing plasmid. IRE1α-oligomerization was induced by the inhibitor of N-linked glycosylation tunicamycin, TM (5 µg/ml) for 4 h. Scale bar, 100µm. In quantification of MANF effect on IRE1α oligomerization upon ER stress the normalized number of IRE1α-GFP foci stands for the number of IRE1α-GFP clusters to total cell count. Statistical analysis was performed using one-way ANOVA, followed by Holm-Sidak’s multiple comparison test, n=3, ****: p < 0.0001. **b**, Interaction of unlabeled titrated purified IRE1α LD (0-3.39 µM) with labeled through His-tag IRE1α LD (20 nM) is affected in presence of increasing concentration of recombinant purified MANF (10 nM-5 µM). Microscale thermophoresis binding curves, showing mean fraction bound values from n=2-4 individual repeats per binding pair ±SEM, Kd values±error estimations are indicated. **c**, The overexpression of MANF K96A and MANF R155A mutants is not affecting IRE1α oligomerization upon ER stress, induced by tunicamycin, TM (5 mg/ml) for 4 hours. Number of IRE1α -GFP foci to total cell count is indicated. Statistical analysis was performed using one-way ANOVA, followed by Holm-Sidak’s multiple comparison test, n=3, **: p < 0.01, ****: p < 0.0001. **e**, Labeled through His-tag luminal domain of IRE1α (20nM) is not interacting with unlabeled titrated recombinant purified MANF K96A mutant protein (0-4.6 µM), while its affinity to MANF K128AC130S protein (0-4.6 µM) is the same as to wtMANF protein (0-4.6 µM), as shown using microscale thermophoresis (MST). Microscale thermophoresis binding curves, showing mean fraction bond values from n=3-5 individual repeats per binding pair ±SEM, Kd values±error estimations are indicated. **f**, Interaction of unlabeled titrated purified IRE1α LD (0-3.39 µM) with labeled through His-tag IRE1α LD (20 nM) is not affected affected in presence of 1µM recombinant purified MANF K96A mutant protein. Microscale thermophoresis binding curves, showing mean fraction bound values from n=3-4 individual repeats per binding pair ±SEM, Kd values±error estimations are indicated.

We further tested MANF effect on IRE1α oligomerization using MST and purified MANF and IRE1α LD proteins. We tested the interaction of unlabeled titrated IRE1α LD with fluorescently labeled IRE1α LD and showed that IRE1α LD is interacting with labeled IRE1α LD with extremely high affinity K_d_=2.7±1.6nM (Fig. 5b). We further analysed whether MANF at different concentrations (10nM-5µM) affects IRE1α LD binding to labeled IRE1α LD, and found that in presence of 100nM MANF the affinity of IRE1α LD to IRE1α LD binding drops to K_d_=40±29nM and in presence of 1µM and 5µM of MANF interaction IRE1α LD-IRE1α LD is abolished (Fig. 5b). This finding confirmed the ability of MANF to decrease IRE1α oligomerization.

### MANF mutant deficient for IRE1α binding does not affect IRE1α oligomerization

To test whether MANF mutants putatively deficient for IRE1α binding have a similar effect on the oligomerization status of IRE1α, we performed transient 48-hour overexpression of MANF mutants in the same way as we did for wtMANF. Two MANF mutants MANF K96A and MANF R155A did not affect the level of oligomerization of IRE1α (Fig. 5c). MANF cysteine loop mutants MANF C130S and MANF K128A C130S decreased IRE1α oligomerization similarly to wtMANF, suggesting that cysteine loop of MANF is not involved in MANF binding to IRE1α.

We expressed and purified recombinant proteins of putatively deficient for IRE1α binding MANF mutants MANF K96A and MANF K128AC130S in CHO cells (Supplementary Fig. 5d). MANF K96A was chosen due to more pronounced difference from wtMANF in IRE1α oligomerization assay as compared with other mutants and MANF K128A C130S was chosen to test also the mutant from other putative binding region predicted by computation modeling. We tested the interactions of purified MANF mutant proteins with fluorescently labeled IRE1α LD using MST. We found that MANF K96A was not interacting with IRE1α LD, while the affinity of MANF K128A C130S to IRE1α LD was not compromised, it was interacting with IRE1α LD with the similar affinity to that of wtMANF with K_d_=55±35 nM (Fig. 5e). We then decided to test, whether MANF K96A is affecting IRE1α dimerization similarly to wtMANF using MST. We tested the interaction of unlabeled titrated IRE1α LD with fluorescently labeled IRE1α LD in presence of 1µM MANF K96A and found that in presence of MANF mutant IRE1α LD is still binding IRE1α LD K_d_=7.3±4.2nM (Fig. 5f). This finding suggests that the direct binding of MANF to IRE1α LD is required for the MANF ability to decrease IRE1α oligomerization.

### MANF is reducing IRE1α phosphorylation upon ER stress

Since MANF decreased IRE1α-oligomerization, we tested the effect of MANF on IRE1α phosphorylation upon ER stress. Phosphorylation of Ser724 in activation loop of IRE1α is believed to be crucial for the activation of IRE1α and triggering of XBP1 splicing (Prischi et al., 2014). We assessed the level of pSer724-IRE1α upon tunicamycin-induced ER stress in transfected/transduced with MANF or MANF K96A HEK293 cells and IRE1α knockout mouse embryonic fibroblasts reconstituted with IRE1α-HA. We showed that in HEK293 cells MANF transfection decreased pSer724 IRE1α after 240 minute-ER stress (Fig. 6x). MANF treatment of IRE1α knockout mouse embryonic fibroblasts reconstituted with IRE1α-HA also reduced pSer724-IRE1α upon 240 minute-(TM)-induced ER stress and 24-hour amino acid starvation (Fig. 6a, b).

**Fig. 6.**
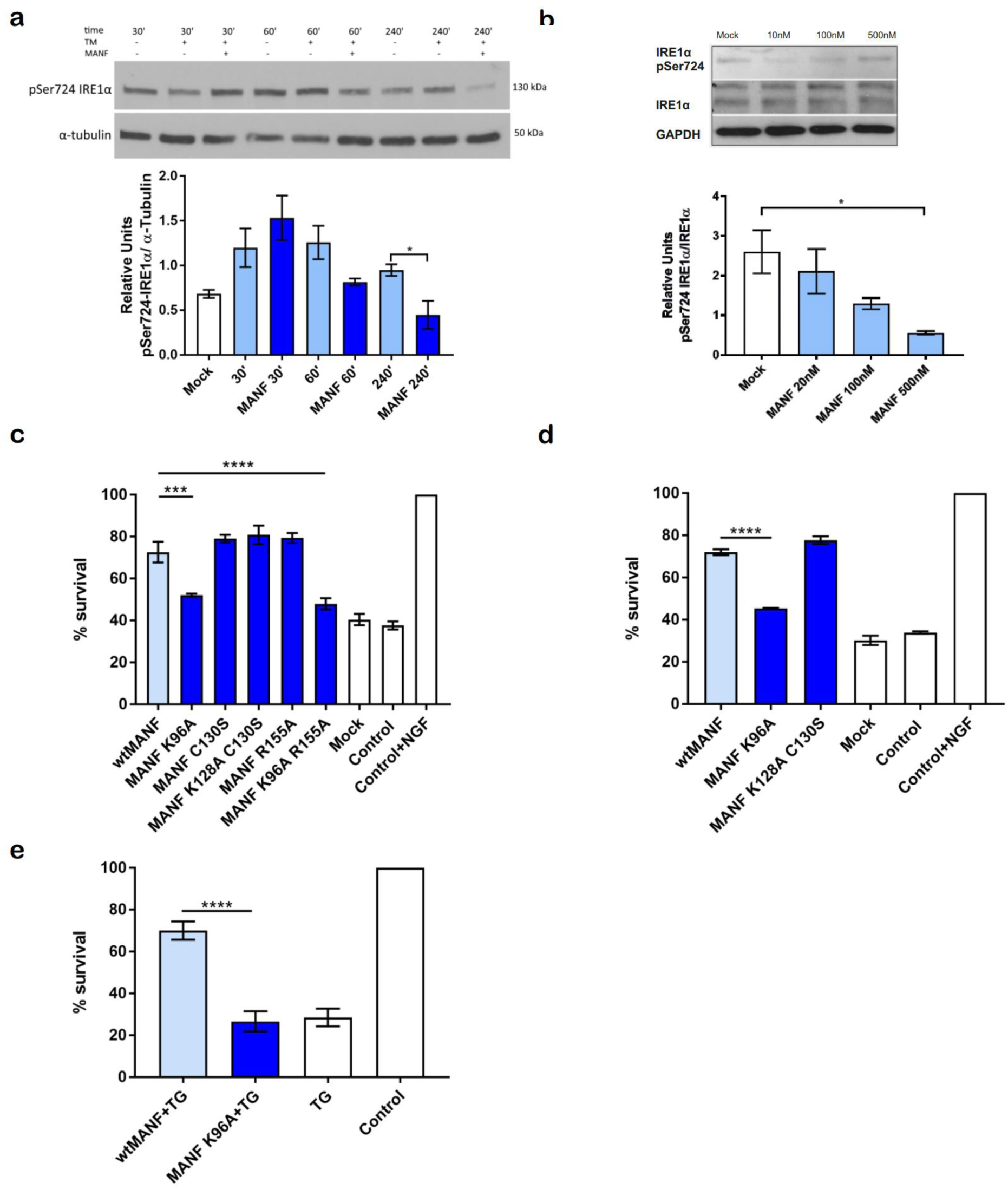
MANF is decreasing phosphorylation of IRE1α. MANF-IRE1α interaction is crucial for the survival of mouse sympathetic and dopamine neurons in ER stress. **a**, Representative image and quantification of pSer724 IRE1α in IRE1α-HA-MEFs treated 4 hours with tunicamycin 1µg/µl, followed by 30, 60 and 240 minutes of treatment with human MANF (50nM), pSer724 IRE1α is normalized to α-tubulin level, the mean±SEM values for n=3 independent experiments are indicated **b**, Representative image and quantification of pSer724 IRE1α in IRE1α-HA-MEFs deprived from amino acids for 24 hours, followed by 24 hour treatment with human recombinant MANF (20, 100 and 500nM), pSer724 IRE1α is normalized to IRE1α level, the mean±SEM values for n=2 independent experiments are indicated. **c**, Microinjections of MANF K96A and MANF K96A R155A MANF mutant plasmids to SCG neurons are not rescuing them from TM-induced apoptosis. **d**, Microinjections of recombinant purified MANF K96A mutant protein to SCG neurons are not rescuing them from TM-induced apoptosis. Statistical analysis was performed using one-way ANOVA, followed by Holm-Sidak’s multiple comparison test, n=3, *: p < 0.05, ***: p < 0.001, ****: p < 0.0001. **e**, MANF K96A is not protecting dopamine (DA) neurons from ER stressed induced apoptosis. DA were cultured 5-7 days *in vitro*, ER stress was induced by treatment with 200 nM thapsigargin (TG). MANF (100ng/ml) or MANF K96A (100ng/ml) were added to the cultures at the same time as TG. Statistical analysis was performed using one-way ANOVA, followed by Holm-Sidak’s multiple comparison test, n=4, ****: p < 0.0001.

### MANF-IRE1α interaction is crucial for the survival of mouse sympathetic in ER stress

MANF, both overexpressed from plasmid or microinjected as a protein have been already shown to be protective against tropoisomerase II inhibitor etoposide, protein kinase inhibitor staurosporine- and nerve growth factor (NGF) deprivation-induced apoptosis in mouse sympathetic SCG neurons (Hellman et al., 2011, Mätlik et al., 2015). Recently, we have also showed that MANF rescues sympathetic neurons from TM-induced apoptosis (Eesmaa et al., under review). Microinjection of wtMANF construct significantly increased the survival of TM-treated SCG neurons: by 35% as compared to empty vector (Fig. 6c). These findings are further supported by our recent study, where IRE1α kinase inhibitor KIRA6 and IRE1α RNase inhibitor 4µ8C abolish the prosurvival effect of MANF on dopamine neurons and sympathetic SCG neurons *in vitro* (Eesmaa et al., under review)

To test whether MANF mutants have similar to wtMANF anti-apoptotic activity, we tested them, first, by microinjecting the respective expression plasmids or mutant proteins into mouse SCG neurons. We found that the mutation at Lys96 (MANF K96A) resulted in the abolishment of pro-survival activity of MANF, double mutant of Lys96 and Arg155 (MANF K96A R155A) was not biologically active either (Fig. 6c). Interestingly, Arg155 mutant (MANF R155A) and both cysteine loop mutants (MANF C130S and MANF K128A C130S) were protecting mouse SCG neurons from TM-induced apoptosis as efficiently as wtMANF.

We then tested the IRE1α LD binding deficient MANF K96A and IRE1α LD-binding MANF K128A C130S mutants by microinjecting the respective mutant proteins to neurons in mouse SCG-neuron survival assay. We found that mutant MANF K96A protein was not able to protect SCG-neurons from TM-induced apoptosis, while the other mutant protein MANF K128A C130S exhibited the same anti-apoptotic activity as wtMANF (Fig. 6d). This result confirms that in SCG-neurons MANF exerts its cytoprotective effect against ER stress through the direct interaction with IRE1α.

Before performing *in vivo* experiment we tested the ability of IRE1α binding-deficient MANF K96A to support the survival of dopamine neurons in culture upon ER stress induced with SERCA-pump inhibitor thapsigargin (TG). We showed that MANF K96A was not able to protect dopamine neurons from TG-induced apoptosis (Fig. 6e).

### MANF mutant unable to bind to IRE1α cannot improve motor behavior in rat 6-OHDA model of Parkinson’s disease *in vivo*

To test whether MANF-IRE1α interaction is important for the neurorestorative activity of MANF *in vivo*, we went on to investigate the effects of wtMANF and MANF K96A mutant proteins in rat 6-OHDA model of Parkinson’s disease (PD), described previously (Lindholm et al., 2007; Voutilainen et al., 2009). Since wtMANF was demonstrated to restore motor behavior after lesioning *in vivo* through the protection of tyrosine hydroxylase (TH)-positive cells in the substantia nigra pars compacta (SNpc), we investigated in current study the behavioral effect of deficient for IRE1α-binding MANF K96A.

The animals were injected with wtMANF, MANF K96A or vehicle intrastriatally two weeks after 6-OHDA lesioning (Supplementary Fig. 7b). Single intrastriatal wtMANF injection reduced ipsilateral turning behavior as compared to vehicle-treated rats with maximal effect at 12 weeks after lesioning (Fig. 7a). MANF K96A injection had no effect on the rotational behavior, confirming that MANF-IRE1α interaction is crucial for restoring motor behavior in the animal model of PD.

**Fig. 7.**
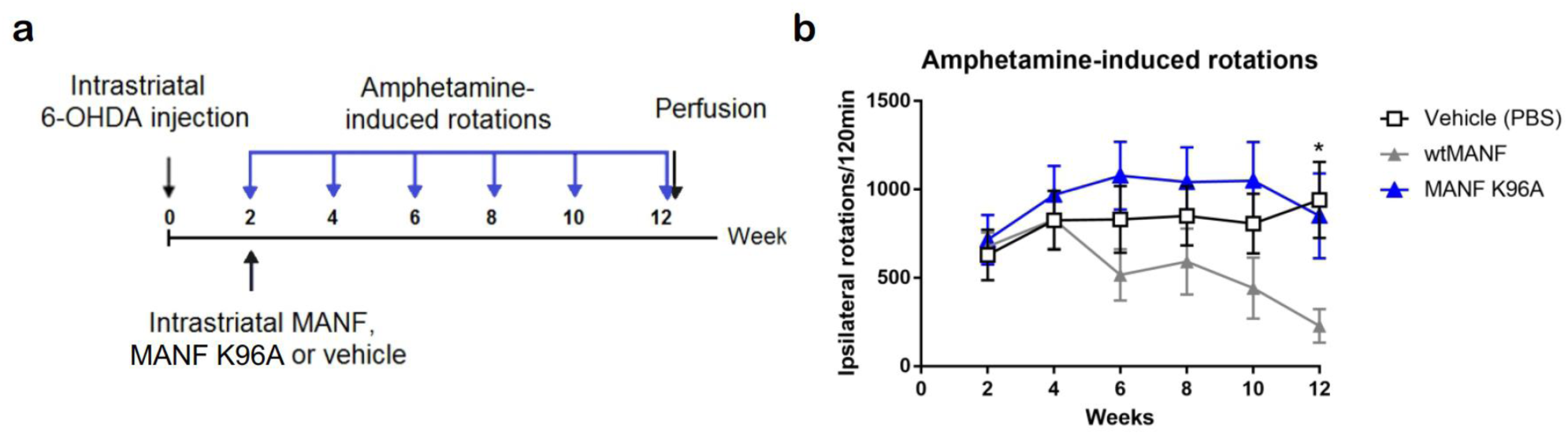
MANF mutant deficient for IRE1α binding does not affect rotational behavior in 6-OHDA model of Parkinson’s disease *in vivo*. **a**, Experimental paradigm for the study. **b**, Amphetamine-induced rotations. Vehicle-treated rats show robust turning behaviour. Single intrastriatal wtMANF injection reduces turning behaviour as compared to Vehicle-treated rats. P=0.0146, ANOVA.

## Discussion

Inositol-requiring enzyme 1α (IRE1α) is the most evolutionary conserved and well studied UPR sensor. However, the mechanisms of its activation and regulation still remain poorly understood. According to classical view the activation of IRE1α and two other UPR sensors, PERK and ATF6 is triggered by the dissociation of major ER chaperone BiP from their luminal domains (Walter and Ron, 2011). Recently the activation of IRE1α has been shown to be triggered by misfolded proteins (Karagöz et al., 2017) and the chaperone Hsp47 (Sepulveda et al., 2018), while protein disulfide isomerase A6 (PDIA6) has been shown to inhibit IRE1α after BiP dissociation from IRE1α LD (Eletto et al., 2014). Similarly to IRE1α, PERK can be also activated by misfolded proteins and other chaperone protein disulfide isomerase A1 (PDIA1) (Wang et al., 2018; Kranz et al., 2017). Due to the physiological importance in keeping cells alive by maintaining cell survival under stress conditions most likely the regulation of IRE1α activation as well as its inhibition under chronic ER stress are complex processes, involving not only BiP, but also other proteins.

Here we show MANF as a major IRE1α interactor and regulator-inhibitor of its activation in ER stress conditions. To date MANF has been shown to alleviate ER stress in various *in vitro* and *in vivo* models (Mizobuchi et al., 2007; Apostolou et al., 2008; Tadimalla et al., 2008; Hellmann et al., 2011; Lindahl et al., 2014; Voutilainen et al., 2017; Danilova et al., 2019; Pakarinen et al. 2020), but the exact mechanism underlying its cytoprotective effects had not been shown before. After MANF has been found to be involved in modulation of innate immunity, the number of studies on MANF dramatically increased (Neves at al., 2016). Despite the growing interest in MANF, only a few papers addressed the mechanism of its action. Binding of MANF to lipid sulfatide has been shown to be promoting cellular uptake of MANF and cytoprotection against hypoxia-induced cell death (Bai at al., 2018). MANF was shown to inhibit nucleotide exchange processes of BiP and prolong the interaction of BiP with misfolded proteins, thereby regulating protein-folding homeostasis (Yan et al., 2019). Our data are not contradicting these studies but rather giving an additional novel understanding of MANF mechanism of action in ER.

The data we present here help to understand why MANF is acting only on ER stressed or injured cells. In the normally functioning cells the major ER chaperone BiP binds with high affinity to luminal domains of IRE1α, PERK and ATF6 and keeps them silent. In ER stress, aggregated or misfolded proteins bind to BiP which is then dissociated from UPR sensors rendering their activation. We show here that MANF binds to LDs of IRE1α, PERK and ATF6 LDs. What is more MANF binds to the same site of IRE1α LD as BiP and therefore MANF binding to IRE1α and possibly to other sensors is possible only, if BiP is dissociated. This means that MANF can bind and regulate IRE1α, and possibly PERK and ATF6 only in highly stressed cells when BiP is dissociated. This also explains, why MANF is not acting on naive healthy cells, because in these cells the MANF binding site in IRE1α LD is occupied and MANF binding is blocked by BiP. According to our results, BiP affinity to IRE1α LD is the lowest and MANF affinity to IRE1α LD is the highest, as compared to the affinities to PERK and ATF6 LDs. These results imply that, in conditions of ER stress BiP dissociates first from IRE1α and thus MANF binds first to IRE1α and only then to PERK and ATF6. MANF-IRE1α binding may therefore be more physiologically significant than binding to other UPR sensors. That is why we focused on IRE1α-MANF interaction; as a continuation of this work the role of interaction of MANF with PERK/ATF6 will be further investigated. Notably, the affinities for mammalian cell produced proteins were higher then for bacterial ones both for BiP and MANF interactions with LDs of UPR sensors likely due to the fact that bacterial cell produced proteins may lack disulfide bonds.

Computational modelling using known 3D structures of MANF, C-MANF and IRE1α LD allowed us to dissect the IRE1α binding site in MANF. We have shown that Lys96 in MANF is involved in the binding with IRE1α and when mutated to alanine, results in abolishment of the effect of MANF on oligomerization of IRE1α, anti-apoptotic activity of MANF in mouse SCG neurons and dopamine neurons and neurorestorative activity of MANF *in vivo* in 6-OHDA model of PD. Lys96 is located in the linker (hinge) region before C-terminal domain of MANF, meaning that more likely the interaction between MANF and IRE1α LD involves in addition to C-MANF also hinge region. In line with that, according to our BIFC data only C-MANF, containing this part of linker region, but not N-MANF was interacting with IRE1α, confirming that more crucial for interaction are amino acids located in the linker region and in the C-terminal domain of MANF. In MST experiments C-MANF showed high affinity to IRE1α LD confirming that binding occurs through C-terminal domain of MANF and hinge region. It suggest that while Lys96 is clearly important for the anti-apoptotic activity of MANF, other amino acids in the C-terminal MANF also play a role in MANF-IRE1α interaction. Further computational studies of the possible structure of MANF-IRE1α/PERK/ATF6 complexes using full-length structure of MANF are of high importance and can help to understand the possible mechanism of interaction of MANF with UPR sensors.

A previous study by Mätlik et al., 2015 showed that cysteine loop mutant MANF C151S (C130S) is not able to rescue ER stressed sensory neurons *in vitro* or in an animal model of stroke. MANF ΔRTDL had a significantly reduced survival effect for sympathetic SCG neurons whereas it was fully active in sensory neurons treated with etoposide. In our systems, we show that MANF C130S reduced IRE1α oligomerization similarly to wtMANF and did not compromise the survival effect of MANF *in vitro*. Thus, the mode of MANF-IRE1α interaction, cell type, intensity of stress and type of cellular stressor may differently affect life and death decisions through IRE1α.

We also demonstrated that MANF decreases IRE1α oligomerization, perhaps due to MANF binding to IRE1α close to the dimerization site of IRE1α. It has been shown before that during ER stress progression MANF expression levels increase and the fraction of IRE1α-bound BiP decreases (Apostolou et al., 2008; Bertolotti et al., 2000). This facilitates MANF binding to IRE1α, which in turn regulates the intensity of IRE1α-mediated UPR response by decreasing or preventing IRE1α-hyperoligomerization, resulting in decreased IRE1α phosphorylation, decreased sXBP1 and decreased apoptosis.

IRE1α can be autophosphorylated at multiple sites (Prischi et al., 2014), and it is not fully clear how phosphorylation at specific phosphorylation sites affects IRE1α activation and downstream signaling. Interestingly, mutation of all three serines in the activation loop of IRE1α does not fully abolish splicing of XBP1, confirming that endoribonuclease activity is not fully dependent on kinase activity of IRE1α or there are other important phosphorylation sites or soluble kinases or phosphatases involved in the regulation of IRE1α activation. In line with this tyrosine kinase c-Abl was shown to increase IRE1α phosphorylation and endoribonuclease activity in chronic ER stress, leading to apoptosis (Morita et al., 2017). MANF is decreasing IRE1α phosphorylation at Ser724 most likely through stabilizing of monomeric conformation of IRE1α and thereby prevention of IRE1α dimerization and autophosphorylation. Alternatively, MANF binding can stabilize IRE1α in conformation favoring the recruitment of soluble kinase or phosphatase affecting IRE1α phosphorylation at Ser724 and possibly at other phosphorylation sites.

Spliced X-box-binding protein 1 (sXBP1) is generally linked to activation of prosurvival mechanisms and restoration of homeostasis upon ER stress (Lee et al., 2003). However, in some cases high level of sXBP1 was reported to result in negative consequences. It was shown to facilitate release of pro-inflammatory extracellular vesicles upon lipotoxic ER stress in hepatocytes (Kakazu et al., 2016). In lipopolysaccharide-induced ER stress *in vivo* sXBP1 was shown to induce acetyltransferase P300 and impair insulin signaling (Cao et al., 2017). We showed that MANF downregulates sXBP1 in HEK293 cells in this study and in dopamine neurons (Eesmaa et al., under review). However, we favor the hypothesis that anti-apoptotic action of MANF is realized not through the downregulation of sXBP1, but rather through prevention of activation of pro-apoptotic pathways, known to be triggered by hyperactivated IRE1α.

Upon chronical severe unresolved ER stress hyperactivated and hyperoligomerized IRE1α is known to recruit TRAF2 and ASK1 and trigger JNK/MAPK-mediated apoptosis and transcription of pro-inflammatory genes (Nishitoh et al., 2002, Brozzi & Eizirik, 2016). IRE1α-hyperoligomerization has been also shown to upregulate thioredoxin interacting protein (TXNIP), activating NLRP3 inflammasome and promoting apoptosis (Morita et al., 2017). Prosurvival action of MANF through prevention of IRE1α hyperoligomerization can be also due to the inhibition of canonical NF-kB pathway shown to be triggered by IRE1α activation through the degradation of IkB by IRE1α upon ER stress (Kaneko et al., 2003; Hu et al., 2006). In line with this MANF has been shown to reduce NF-kB pathway activation in beta cells (Hakonen et al., 2018) and in HEK293T cells (Chen et al., 2015).

Based on our findings and previous studies on IRE1α-phosphorylation-hyperoligomerization we propose the following putative mechanism of MANF signaling through IRE1α (Fig. 8). We suggest that MANF represents a ‘second wave’ of UPR regulation-inhibition, following BiP dissociation from IRE1α. MANF via its C-terminal domain and linker region is binding IRE1α close to its dimerization surface and prevents IRE1α dimerization-oligomerization and hyperactivation upon chronic ER stress. That leads to decrease in phosphorylation of IRE1α, reduced sXBP1 formation and prevents triggering of IRE1α hyperactivation-induced pro-apoptotic mechanisms and therefore increases cell survival. In chronic ER stress MANF eventually also binds to PERK an ATF6 and through these interactions can regulate them as well. In MANF KO mice first IRE1α, but later all three UPR pathways are activated. We showed earlier that MANF can also attenuate the downstream signalling of PERK and ATF6 pathways (Eesmaa et al., under review) and, perhaps, the specificity of attenuation of UPR sensors is determined by order and degree of BiP dissociation from UPR sensors, correlating with the severity of ER stress. MANF might be also involved in the crosstalk between UPR sensors.

**Fig. 8.**
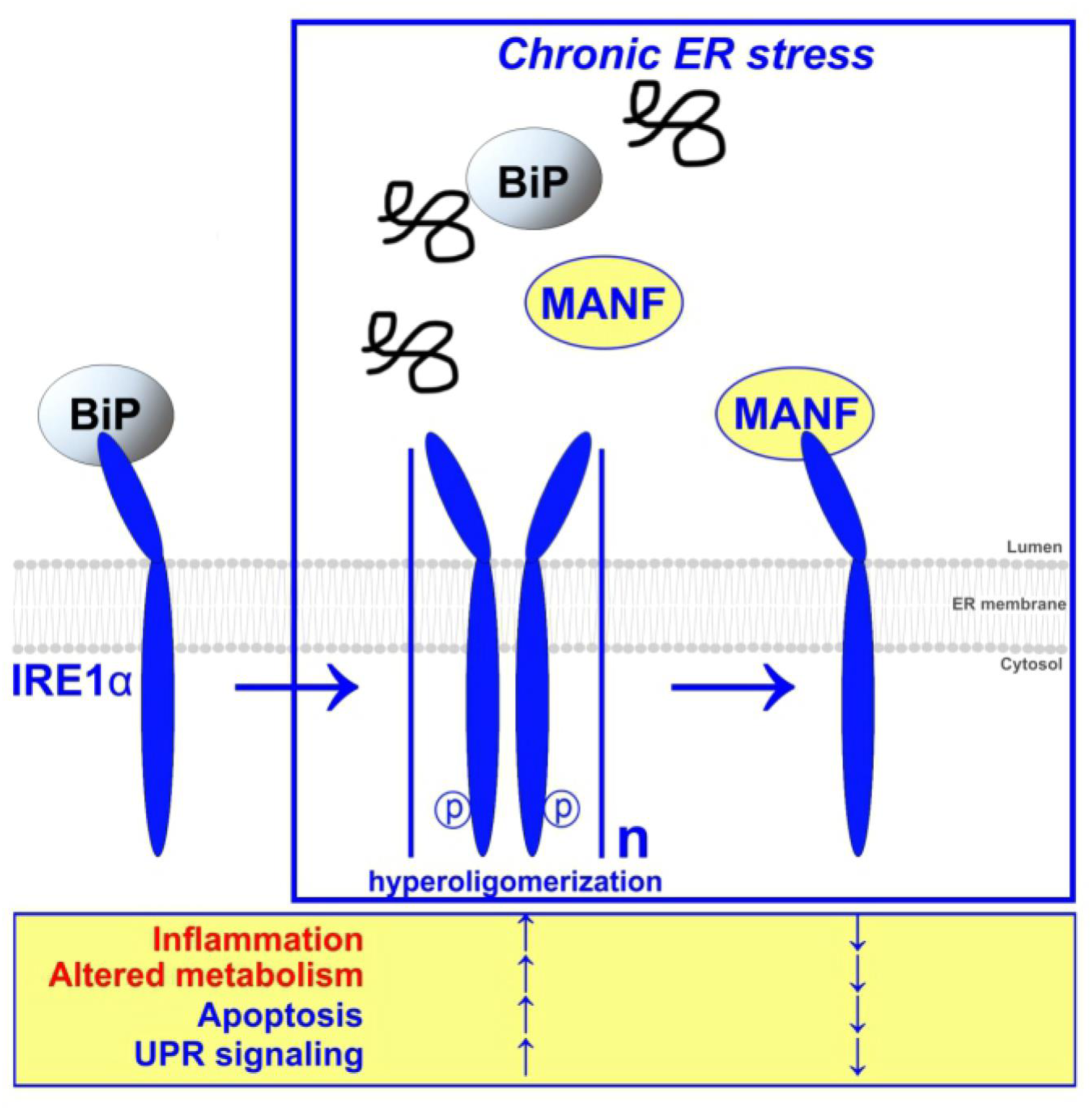
Putative mechanism of MANF signaling through IRE1α. Upon ER stress when BiP dissociates from IRE1α luminal domain, MANF directly binds to IRE1α preventing IRE1α hyperoligomerization and decreasing IRE1α phosphorylation, resulting in neuroprotective and neurorestorative effect both *in vitro* and *in vivo*.

Our results are important for the development of new strategies to treat different neurodegenerative and UPR associated diseases, such as diabetes. Specific inhibitor of IRE1α signaling KIRA8, have already shown to have anti-diabetic effect (Ghosh et al., 2014; Morita et al., 2017). Another attenuator of IRE1α signaling c-Abl inhibitor Imatinib (Gleevec) was shown to restore cognitive function and have neuroprotective potential in LPS-induced inflammatory mouse model (Weintraub et al., 2013) and also reduces brain injury after traumatic brain injury (Su et al., 2015). Considering that according to our results MANF can act similarly, but interacts with all three UPR sensors (IRE1α, PERK, ATF6), MANF can be much more potent as neuroprotective and anti-diabetic agent. In future *in vitro* screening of small-molecule compounds, mimicking MANF action on IRE1α and other UPR sensors can lead to the development of new drugs.

## Materials and Methods

### Cell lines

HEK293 cells for bimolecular fluorescence complementation assay (BiFC) experiments were grown in Dulbecco’s modified Eagle’s medium DMEM supplemented with 10% fetal bovine serum and 50ug/ml normocin (ant-nr-2, Invivogen). HEK293 cells and other cell lines used were grown at 37°C and 5% CO_2_.

Flp-In™ T-REx™ 293 cell line (Invitrogen) containing a single stably integrated FRT site and expressing Tet repressor were used for generation of inducible cell lines, expressing IRE1α-HA, BiP-HA or GFP-HA. The medium composition was the same as for HEK293 cells.

Flp-InTM-CHO cells (R75807, Thermo Fisher Scientific) were grown in growth media consisting of Ham’s F12 nutrient mix (21765029, Thermo Fisher Scientific), 10% FBS (10270106, Gibco), 2mM GlutaMAX (35050061, Thermo Fisher Scientific) and normocin (ant-nr-2, Invivogen). We used the Flp-InTM-CHO to generate CHO-derived stable transgenic cell lines overexpressing either MANF or its mutants from a transcriptionally active locus. For this, the respective pcDNA5/FRT/TO pre-SH-MANFwt or mutant constructs were cotransfected with the Flp-recombinase expressing pOG44 plasmid in a 1:9 ration using JetPEI (101-10N, Polyplus Transfections) transfection reagent. The selection was started 48 hours after transfection using growth media supplemented with 500µg/ml Hygromycin Gold (ant-hg-1, Invivogen). Selection media was changed every 3-4 days until confluent colonies of stable transgenic cells had formed and cells were ready to be split.

Mouse embryonic fibroblasts (MEFs) cells were grown in Dulbecco’s modified Eagle’s medium DMEM (12-614F, Lonza), supplemented with 5% fetal bovine serum and non-essential amino acids at 37°C and 5% CO_2_.

### Reagents and proteins

The inducers of ER stress thapsigargin (T7459, ThermoFisher Scientific) and tunicamycin (ab120296, Abcam) and IRE1α inhibitors KIRA6 (HY-19708, MedChemExpress) and 4µ8C (14003-96-4, Cayman Chemical) were used.

Luminal domains (LD) of three UPR sensors, human IRE1α, PERK and ATF6 and human MANF were expressed and purified in *CHO* cells by Icosagen Ltd (Tartu, Estonia). Human C-MANF was expressed and purified from *E*.*coli* cells (Hellmann et al., 2011) or synthesized chemically by Apeptide Ltd (Shanghai, China). Human recombinant GRP78 (BiP) (SMB-SPR-119A, StressMarq Biosciences Inc.) was used.

### Microscale thermophoresis (MST)

The experiments have been performed using Monolith NT.115 instrument (NanoTemper Technologies GmbH, Germany). Recombinant proteins were labeled through His-tag using Monolith His-Tag Labeling Kit RED-tris-NTA (MO-L008). The concentration of labelled proteins (targets) was 20nM for all the experiments and different starting concentrations of the ligands have been used. Experiments were performed in a buffer containing 10mM Na-phosphate buffer, pH 7.4, 1mM MgCl_2_, 3mM KCl, 150mM NaCl, 0.05% Tween-20. The measurements were done in premium coated capillaries (NanoTemper Technologies GmbH, MO-K025) using red LED source, power set at 100% and medium MST power at 25°C. Each data point represents mean ΔFnorm values from n=2-4 independent experiments per binding pair ±S.D, Kd values±error estimations are indicated. Data analysis was performed using MO.Affinity Analysis v2.3 and GraphPad Prism 7 software.

#### Binding assay on nickel-coated plates

Nickel-coated plates (15442, Pierce) were blocked with 1% Casein PBS-T for 1 hour at RT on shaker, Bip-His+MANF (positive control), IRE1α LD-His+MANF and GFRα1-His+MANF (negative control) 1:1 mixtures were prepared in the buffer, that was used for microscale thermophoresis experiments. MANF+buffer mixture (1:1) was included to measure the background absorbance. The protein mixtures were vortexed and incubated at RT for 1 hour. After the incubation protein solutions were pipetted on the plates and incubated on the plates for 1 hour at RT. The plates were washed 3 times with 0.05% Tween-20 (P2287, Sigma-Aldrich) in PBS. After washing the detecting antibody HRP-linked mouse anti-human MANF, clone 4E12 (Icosagen) in blocking buffer was added and the plates were incubated overnight at +4°C on shaker. Next day the plates were washed 3 times and the color development was performed using Duoset ELISA Development System (DY999, R&D Systems) according to the manufacturer’s instructions. The absorbance at 450 and 540nm was measured using plate reader (VICTOR3, Perkin Elmer). The background absorbance was subtracted. Data analysis was performed using GraphPad Prism 7 software

### Analysis of purity and glycosylation of luminal domains of UPR sensors

For analysis of protein N-glycosylation PNGase F (P0704S, New England Biolabs) was used and the assay was performed according to the manufacturer’s instructions. Glycosylated protein (5µg/well) versus deglycosylated protein was loaded onto mini-PROTEAN precast gels (456-1093, Bio-Rad) and run at 40mA for 1 hour. Coomassie Brilliant Blue G-250 staining was performed according to the standard protocol. Glycosylated mammalian cell produced GDNF protein was used as a positive control.

### Size exclusion chromatography (SEC)

For complex preparation, purified human *CHO* expressed MANF and IRE-1 LD were combined in a molar ratio of 1.25:1 (MANF:IRE1 LD) at a total protein concentration of 0.7 mg/ml in size exclusion chromatography buffer (10 mM MES-NaOH pH 5.5, 150 mM NaCl, 3 mM KCl, 1 mM MgCl_2_, 0.05% TWEEN-20). The complex was incubated for 10 min at 22°C before size exclusion chromatography using a Superdex 200 Increase 3.2/300 column (GE Healthcare) at 22°C. Individual components were also run similarly. Selected fractions were analysed by Western blotting.

### Protein – protein docking

The X-ray diffraction crystal structure of the human IRE1-alpha luminal domain (IRE1α, PDB ID: 2HZ6) with resolution 3.1 Å (Zhou et al., 2006) and the structure with the least restraint violations from the NMR solution structure of the C-terminal domain of mesencephalic astrocyte-derived neurotrophic factor (MANF, PDB ID: 2KVE) (Hellman et al., 2011) were used for protein–protein docking. The binding poses of the IRE1α - MANF complexes were predicted by docking calculations using Schrödinger LLC BioLuminate software (Schrödinger 2018a, Schrödinger 2018b, etc.). Before molecular docking, the 3D structure of protein molecules was optimized using the Protein Preparation Wizard (OPLS_2005 force field) in the Schrödinger LLC Maestro software (Schrödinger 2018a, Schrödinger 2018b, etc.). The protein–protein docking was carried out using PIPER procedure, which proceeds with a rigid body global search based on the Fast Fourier Transform (FFT) correlation approach (Kozakov et al., 2006). The PIPER procedure performs exhaustive evaluation of an energy function in discretized 6D space of mutual orientations of two proteins. The structures corresponding to different mutual orientations of the proteins were ordered according to the scoring function that is given as the sum of terms representing shape complementarity, electrostatic, and desolvation contributions The top 1000 structures were subsequently clustered using the pairwise root mean square deviation (RMSD) as the distance measure between two proteins in the complex within a fixed clustering radius 9 Å (Kozakov et al., 2005). The selected structures from30 largest clusters were refined by a SDU (Semi-Definite programming based Underestimation) medium-range optimization method (Paschalidis et al, 2007). The analysis of the protein–protein interactions of the final 30 top configurations was performed by using AutoDock Tools software (Morris et al., 2009).

### Generation of MANF mutant plasmids and recombinant proteins

pcDNA5/FRT/TO pre-SH-MANF K96A, C130S and R155A mutants were generated using site-directed inverse PCR mutagenesis and pcDNA5/FRT/TO pre-SH-MANF as template. The pcDNA5/FRT/TO pre-SH-MANF K128A C130S mutant was generated using pcDNA5/FRT/TO pre-SH-MANF C130S as template.

Recombinant human MANF protein was produced from a CHO-derived cell line using the QMCF technology as has been described before (P-101-100, Icosagen). The MANF K96A and K128A C130S mutant recombinant proteins were produced by Icosagen using the same technology. Briefly, codon-optimized cDNAs were cloned to pQMCF-T expression vectors which were then transiently transfected to CHO-derived protein production cell line. Proteins were captured and purified from the cell culture media using 5ml Q FF followed by 1ml SP HP, buffer was exchanged into PBS pH 7.4 by size exclusion chromatography. Protein purity was verified by SDS-PAGE with Coomassie staining and immunoblotting using rabbit anti-MANF antibody (310-100, Icosagen).

### Expression and secretion of MANF mutant plasmids

CHO cells grown on 6cm plates transfected with pTO expression plasmids. Plasmid DNA (6ug) +12ul of Turbofect per plate. Media changed 24h after transfection to serum-fee media (3ml per 6cm plate). Incubated for 24h more before harvesting cells and collecting media. Each cell pellet was lysed in 400ul of lysis buffer, 100ul of media was set aside before concentrating, and the rest was concentrated from ∼ 3ml to ∼ 100ul using Amicon Ultra-4 centrifugal filters 10K. 20ul of each sample was loaded onto 4-15% gel. Primary ab: m@HA (Abcam) 1:1000 2h. Secondary ab: g@mouse 690 LR (Licor) 1:10000

### Duolink proximity ligation assay (PLA)

The experiments were performed in 96-well format on Flp-in-TREx293 cells, expressing IRE1-HA/BiP-HA/GFP-HA upon doxycycline induction or on CHO cells, stably expressing HA-tagged MANF. 10000 cells per well was plated on pre-coated with Poly-D-Lysine (0.1mg/ml) black Perkin Elmer plates. Cells were fixed with 4% paraformaldehyde for 15 min and afterwards permeabilized/stained with DAPI (D9542, Sigma-Aldrich) in 1xPBS containing 0.05% Triton X-100 for 10 min. Blocking and incubation with antibodies have been performed following Duolink manufacturer’s protocol. Cells were incubated overnight at +4°C with the following primary antibodies: anti-MANF rabbit pAb (Icosagen, 310-100), anti-IRE1α rabbit mAb (CST, 3294), anti-BiP rabbit mAb (CST, 3177), anti-HA mouse mAb (Abcam, ab130275). Incubation with PLUS and MINUS PLA probes have been performed for 1 hour at +37°C. Ligation and amplification was performed according to manufacturer’s instructions. The imaging of 16x sites/well was performed in TexasRed and DAPI channels using MolecularDevices Nano scanner. The analysis and quantification was done using CellProfiler 3.1.5 and CellProfiler Analyst 2.2.1 software.

### Plasmids for BiFC

pCE-BiFC-VC155 (CV) and pCE-BiFC-VN173 (NV) were a gift from Chang-Deng Hu (Addgene plasmids #22020 and #22019). pEZYflag and pEZYmyc-His were a gift from Yu-Zhu Zhang (Addgene plasmids #18700 and #18701). Gateway destination vectors for BiFC for N- and C-terminal tagging with Venus fluorescent protein fragments (pEZY BiFC N NV, pEZY BiFC N CV, pEZY BiFC C NV and pEZY BiFC C CV) were generated by PCR by replacing the flag or myc-His sequences from pEZYflag or pEZYmyc-His with VC155 or VN173 sequences from the respective plasmids.

To generate the MANF Gateway compatible entry vector, pCR3.1 MANF (Hellman et. al., 2010) was used to clone the MANF coding region into pENTR221 vector using Gateway entry clone generation by PCR, following the manufacturer’s instructions (Invitrogen, Thermo Fisher Scientific, USA).

The following Gateway entry clones were from the Genome Biology Unit (GBU) Core Facility (Research Programs Unit, Faculty of Medicine, HiLIFE, University of Helsinki, Biocenter Finland): HSPA5 (Grp78) without stop (DQ895368), JUN without stop (DQ896432), FOS without stop (DQ893444), MAX without stop (JF432558). Shown is the Genbank accession number and the presence or absence of a translation stop-codon to indicate subsequent N- or C-terminal fusion, respectively, with a Venus fragment. The corresponding BiFC expression plasmids were made by LR clonase recombination reaction of pENTR221 constructs into the respective pEZY BiFC destination vector. pDONR223-ERN1 (IRE1) was a gift from William Hahn & David Root (Addgene plasmid # 23491). A stop codon at the end of the IRE1-coding reading frame was added before using that construct to generate a pDONR223 pre-CV-IRE1. For that purpose, inverse PCR was used to linearize the pDONR223 IRE1 with stop construct between the sequences corresponding to the pre-sequence and the mature IRE1. A C-Venus insert with a GS-linker was amplified from a pEZY BiFC pre-CV containing construct and used in a ligation reaction with the linearized pDONR223 IRE1 stop construct to generate pDONR223 pre-CV-IRE1. The latter was then used as an entry clone in an LR reaction to recombine the pre-CV-IRE1 sequence into the pEZY BiFC myc-His destination vector.

pEZY BiFC Grp78 NBD-NV and Grp78 SBD-NV were made using Grp78 NBD and SBD specific primers, inverse PCR and pEZY BiFC Grp78-NV as a template.

pENTR221 pre-N-Venus MANF was generated by amplifying the sequence corresponding to VN173 from the respective BiFC destination vectors and inserting it between the sequences coding for signal peptide (pre) and mature regions of human pENTR221 MANF. The corresponding BiFC expression plasmids (pEZY BiFC pre-NV-MANF) was made by LR clonase recombination reaction of pENTR221 pre-NV-MANF into pEZY Myc-His destination vector.

pEZY BiFC pre-NV-N-MANF and pre-NV-C-MANF constructs were generated using inverse PCR reactions the N-MANF or C-MANF specific primers, respectively and the pEZY BiFC pre-NV-MANF as a template.

### Bimolecular fluorescence complementation assay (BiFC)

HEK293 cells plated on covered with Poly-D-Lysine (P0899, Sigma-Aldrich) coverslips were co-transfected with pEZY BIFC N-Venus and C-Venus plasmids 48 hours after plating. Transfection with jetPEI transfection reagent (101, Polyplus-transfecton) was performed according to the manufacturer’s protocol. 20 hours after transfection the cells were fixed with 4% PFA, washed with PBS and permeabilized using 0.1% Triton X-100 in PBS (PBS-T). For nuclear and endoplasmic reticulum staining we used ER-ID® Red assay kit (ENZ-51026-K500, Enzo Life Sciences), containing Hoechst 33342 nuclear stain and ER-ID® Red detection reagent. ProLong™ Diamond Antifade Mountant (P36965, ThermoFisher Scientific) was used for mounting of coverslips. The imaging was performed using Leica SP8 STED confocal microscope, 63x glycerol immersion objective and Leica Application Suite X (LASX) software. Image analysis and processing (including brightness&contrast adjustment, same for all images) was done using CorelDRAW 2018.

### IRE1α oligomerization assay

TREx-293IRE1α-3FGHGFP cells (Li et. al., 2010) were plated 5000 cells/well on pre-coated with Poly-D-Lysine (0.1mg/ml) black Perkin Elmer plates in DMEM with 10% FBS and 100ug/ml Normocin. Next day the cells were transiently transfected with pTO-pre-SH-MANF-GW-FRT (MANF mutants) or pTO-SH-GW-FRT as a control vector, 100ng of plasmid/well for 24 hours using PEI Transfection Reagent (1ug/ul in 1x PBS pH 4.5; 4:1 v/w ratio of PEI:DNA). After transfection, IRE1α-GFP expression was induced with doxycycline (1ug/mL) treatment for 24 hours. ER stress was induced by treating the cells with the inhibitor of N-linked glycosylation tunicamycin (TM), 5ug/ml for 4 hours. After treatment, cells were fixed with 4% paraformaldehyde for 20 min and stained with DAPI (D9542, Sigma-Aldrich) in 1xPBS for 10 min. Imaging (16 sites/well) was performed using MolecularDevices Nano scanner. Three independent experiments have been analysed and quantified using CellProfiler 3.1.5 and CellProfiler Analyst 2.2.1 software.

### Western blot analysis I

HEK293 cells were plated 250000/well on 12 well plates in DMEM with 10% FBS and 100ug/ml normocin. Next day transient transfection of pTO-pre-SH-MANF-GW-FRT (MANF mutants) or pTO-SH-GW-FRT as a control was performed in the similar way as for IRE1α oligomerization assay, 500ng of plasmid/well. 24 hours after transfection, the cells were starving for 4 hours (DMEM without FBS), and then treated with tunicamycin (TM) 5ug/ml for the times indicated. The lysis was performed in RIPA buffer (1% SDS, 10 mM Tris pH 8.0, containing protease and phosphatase inhibitor cocktail tablets (Roche)). The concentrations of total protein in cell lysates were measured using NanoDrop 2000 spectrophotometer. 20ug/well of total protein was loaded onto Bio-Rad mini-PROTEAN precast gels followed by the transfer onto nitrocellulose membrane for conventional western blotting or SuperSep™ Phos-tag™ (50µmol/L) Zn^2+^ precast gels (FUJIFILM Wako, 198-17981) followed by the transfer onto polyvinylidene difluoride (PVDF) membrane for the detection of phosphorylated IRE1α using the Phos-tag™ assay. The transfer have been done for 1hour on ice at RT. Pre-treatment of the membranes with EDTA for Phos-tag™ assay was performed according to the manufacturer’s instructions. The membranes were blocked in 5% BSA TBS-T (or 5% milk TBS-T) for 1 hour at room temperature and then incubated with primary antibodies overnight at +4°C. The following primary antibodies were used: anti-IRE1α (CST, 3294), IRE1α pSer 724 (NovusBio, NB100-2323), GAPDH (EMD Millipore, MAB 374). Peroxidase-coupled secondary antibodies and the enhanced chemiluminescence (ECL) detection system have been used for western blot development.

### Western blot analysis II

MEFs cells were plated 250000/well on 12 well plates in DMEM with 5% FBS and non-essential amino acids. Next day the cells were treated with tunicamycin (TM) 500ng/ml for 4 hours, followed by treatment with exogenous human recombinant MANF (50nM) for 30, 60 and 240 minutes. The lysis was performed in RIPA buffer, containing protease and phosphatase inhibitor cocktail tablets (Roche)). The concentrations of total protein in cell lysates were measured using NanoDrop 2000 spectrophotometer. 20ug/well of total protein was loaded onto Bio-Rad mini-PROTEAN precast gels followed by the transfer onto nitrocellulose membrane. The transfer have been done for 1 hour on ice at RT. The membranes were blocked in 5% BSA TBS-T (or 5% milk TBS-T) for 1 hour at room temperature and then incubated with primary antibodies overnight at +4°C. The following primary antibodies were used: anti-IRE1α (CST, 3294), IRE1α pSer 724 (NovusBio, NB100-2323), α-Tubulin (Sigma-Aldrich, T9026). Peroxidase-coupled secondary antibodies and the enhanced chemiluminescence (ECL) detection system have been used for western blot development.

### Neuronal culture and microinjection

Culture of mouse superior cervical ganglion sympathetic neurons and microinjection of these neurons was performed as described earlier (Yu et al., 2003). Briefly, the neurons of postnatal day 1–2 NMRI strain mice were grown 6 DIV on polyornithine-laminin (P3655 and CC095, Sigma-Aldrich)–coated dishes or glass coverslips with 30 ng/ml of 2.5 S mouse NGF (G5141, Promega) in the Neurobasal medium containing B27 supplement (17504044, Invitrogen). The nuclei were then microinjected with the expression plasmid for full-length MANF (pTO-pre-SH-MANF) together with a reporter plasmid for enhanced green fluorescent protein (EGFP), at concentration of 10 ng/ul in each experiment. For protein microinjection, recombinant full length MANF protein (P-101-100, Icosagen) in PBS at 200ng/ul was microinjected directly into the cytoplasm together with fluorescent reporter Dextran Texas Red (MW 70000 Da) (D1864, Invitrogen, Molecular Probes) that facilitates identification of successfully injected neurons. Next day, tunicamycin (2 µM) (ab120296, Abcam) was added and living fluorescent (EGFP-expressing or Dextran Texas Red-containing) neurons were counted three days later and expressed as percentage of initial living fluorescent neurons counted 2–3 hours after microinjection.

### Immunocytochemistry

The neurons were cultured on glass coverslips and microinjected after 6-7 days in vitro with plasmid encoding for human wtMANF or its mutants. DNA concentration of 10 ng/μl was used. The cells were fixed with 4% PFA at 24 h after microinjection and stained with the following antibodies: rabbit anti-MANF (used in Lindholm et al. 2014), mouse anti-PDI (ADI-SPA-891-F, Enzo Life Sciences), mouse anti-GM130 (610823, BD Biosciences), goat anti-GRP78 (sc-1051, Santa Cruz Biotechnology Inc.), Alexa Fluor 488 goat anti-rabbit IgG (H+L) (A-11008, Invitrogen) and Alexa Fluor 568 goat anti-mouse IgG (H+L) (A-11004, Invitrogen). The nuclei were stained with DAPI (D9542, Sigma-Aldrich). The fluorescent image stacks were acquired using the confocal microscope TCS SP5 equipped with LAS AF 1.82 (Leica Microsystems Inc). The objective was Leica HCX PL APO x63/1.3 GLYC CORR CS (21 °C). The lasers used were DPSS 561 nm/20 mW, OPSL 488 nm/270 mW and diode 405 nm/50 mW, with the beam splitter QD 405/488/561/635. The confocal images were analysed by Imaris 9.2.1 software (Bitplane).

### Primary cultures of midbrain dopaminergic neurons and MANF mutant treatment

The midbrain floors were dissected from the ventral mesencephalic of 13 days old NMRI strain mouse embryos. The tissues were incubated with 0.5% trypsin (103139, MP Biomedical) in HBSS (Ca^2+^/Mg^2+^-free) (14170112, Invitrogen) for 20 min at +37°C, then mechanically dissociated. Cells were plated onto the 96-well plates coated with poly-L-ornithine (Sigma-Aldrich). Equal volumes of cell suspension were plated onto the center of the dish. The cells were grown for 5 days without any neurotrophic. Then, the cells were treated with thapsigargin (20nM) (T7458, Thermo Fisher Scientific) and wtMANF (100ng/ml) or MANF K96A (10ng/ml, 100ng/ml, 1 µg/ml). After three days the neuronal cultures were fixed and stained with anti-Tyrosine Hydroxylase antibody (MAB318, Millipore Bioscience Research Reagents. Images were acquired by CellInsight high-content imaging equipment. Immunopositive neurons were counted by CellProfiler software and the data was analysed by CellProfiler analyst software. The results are expressed as % of cell survival compared non toxin treatment neurons.

### Testing MANF mutant in *in vivo* 6-OHDA model

#### Experimental animals

Male Wistar rats (weight 230-270 g, Envigo, Netherlands) were housed in groups of 3 to 4 under a 12-h light-dark cycle at an ambient temperature of 20–23 °C. Food pellets (Harlan Teklad Global diet, Holland) and tap water were available ad libitum. Experiments were performed according to the 3R principles of EU directive 2010/63/EU on the care and use of experimental animals, as well as local laws and regulations, and were approved by the national Animal Experiment Board of Finland (protocol approval number ESAVI/12830/2020). All experiments were performed in a blinded manner and the rats were assigned to the treatment groups equally based on their rotational score at week 2.

#### 6-OHDA lesioning

6-OHDA injections were done under isoflurane anesthesia essentially as described earlier (Penttinen et al., 2016, Voutilainen et al., 2009; Voutilainen et al., 2011). The animals received unilateral injections totaling 6 µg of 6-OHDA (Sigma Chemical CO, St. Louis, MO, USA; calculated as free base and dissolved in ice-cold saline with 0.02% ascorbic acid) in 3 deposits (2 µg / 1.5 µl each) in the right striatum using coordinates relative to the bregma (A/P + 1.6, L/M + 2.8, D/V−6; A/P 0.0, L/M +4.1, D/V -5.5 and A/P −1.2, L/M +4.5, D/V −5.5) (Paxinos and Watson, 1997). The rats were divided into treatment groups according to their amphetamine-induced rotations on two-week post lesion. After the behavioural tests, the rats were transcardially perfused and their brains were processed for TH immunohistochemistry.

#### Intrastriatal administration of compounds

MANF and mutant MANF were intrastriatally administered to 6-OHDA lesioned rats two weeks after lesioning under isoflurane anesthesia using the same stereotaxic coordinates as with 6-OHDA injections. MANF and mutant MANF were injected in three locations in the striatum in three injections of equal volume. The total injected doses were for MANF and mutant MANF 10 µg. The total injection volume was adjusted to be 2 µl for all compounds.

D-Amphetamine-induced rotational behavior was measured at 2, 4, 6 and 8 weeks post lesion in automatic rotometer bowls (Med Associates, Inc., Georgia, USA) as previously described (Lindholm et al., 2007, Ungerstedt and Arbuthnott, 1970). Following a habituation period of 30 min, a single dose of D-amphetamine (2.5 mg/kg, Division of Pharmaceutical Chemistry, Faculty of Pharmacy, University of Helsinki, Finland) was injected intraperitoneally (i.p.). The rotation sensor recorded complete (360**°**) clockwise and counterclockwise-uninterrupted turns for a period of two hours and ipsilateral rotations were assigned a positive value.

#### Quantification and statistical analysis

GraphPad Prism 7.0 software was used for statistical analysis. Statistical tests and sample sizes are indicated in the figure legends. A p < 0.05 was considered statistically significant.

## Acknowledgements

The study has been supported by Jane and Aatos Erkko Foundation and Sigrid Jusélius Foundation. Lucie Küll, Susanna Wiss, Mari Heikkenen, Carina Gutenbrunner, Iannis Charnay, Karola Meininghaus and are thanked for their technical help. Peter Walter is thanked for sharing the IRE1α-3FGH-GFP construct. We are grateful to Claudio Hetz for sharing the IRE1α^-/-^ MEFs cells and for fruitful discussions. We thank Mikko Airavaara, Andrii Domanskyi and Maria Lindahl for critical comments.

## Author Contributions

VK designed and performed MST, PLA, BiFC, WB, PhosTag-WB, oligomerization experiments, analysed the data, prepared the figures and wrote the manuscript. LY have performed survival studies in SCG and DA neurons, ICC of MANF mutants. LI and MK planned and performed molecular docking experiments. MV designed and performed *in vivo* experiments with MANF mutant. JN performed the analysis of *in vivo* experiments with MANF mutant. AE designed molecular cloning for BIFC experiments and generated MANF mutant constructs. EPK and JH designed and performed gel filtration chromatography and helped in structural analysis. MS supervised the experiments, participated in the designs, in manuscript writing and provided funding. All the authors read and approved the final version of the manuscript.

---

**Supplementary Figure 1.**
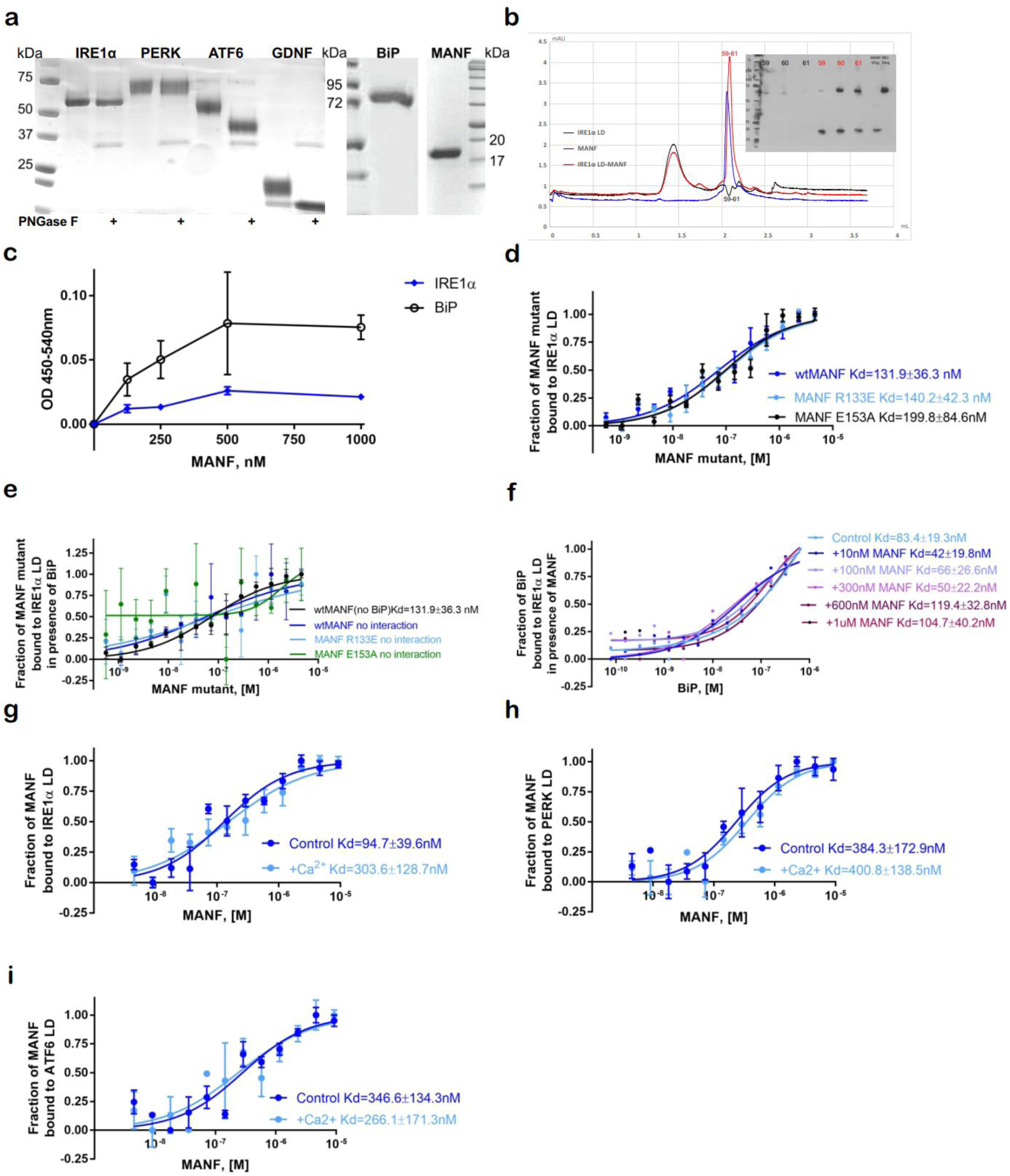
MANF interaction with luminal domains of UPR sensors. **(A)** SDS-PAGE gel of purified from *CHO* cells luminal domains of UPR sensors IRE1α, PERK and ATF6, *E*.*coli* produced human recombinant BiP and *CHO* cells produced human recombinant MANF. Glycosylation of LDs of UPR sensors was tested using PNGase F assay. **(B)** MANF and IRE1α LD form a complex, as shown by SEC, followed by western blotting using MANF and His-tag antibodies. Red curve-IRE1α LD alone, blue curve-MANF alone, black curve-IRE1α LD complex. **(C)** Recombinant purified human MANF protein is interacting with BiP-His and IRE1α LD-His proteins on Pierce™ nickel coated plates. Relative absorbance at 450nm-540nM is indicated. **(D)** Interaction of unlabeled titrated human recombinant *CHO* cells produced deficient for BiP binding MANF E153A and MANF R133E (0-4.6 µM) with Alexa647-labeled through His-tag luminal domains of IRE1α LD (20nM), analyzed using MST. Microscale thermophoresis binding curves, showing mean fraction bond values from n=3 individual repeats per binding pair ±SEM, Kd values±error estimations are indicated. **(E)** Interaction of unlabeled titrated human recombinant *CHO* cells produced deficient for BiP binding MANF E153A and MANF R133E (0-4.6 µM) with Alexa647-labeled through His-tag luminal domains of IRE1α LD (20nM) in presence of 50 nM BiP, analyzed using MST. Microscale thermophoresis binding curves, showing mean fraction bond values from n=3 individual repeats per binding pair ±SEM, Kd values±error estimations are indicated. **(F)** Interaction of unlabeled titrated human recombinant BiP (0-640nM) with Alexa647-labeled through His-tag luminal domain of IRE1α (20nM) in presence of increasing concentrations of human recombinant MANF (10nM-1µM). Microscale thermophoresis binding curves, showing mean fraction bond values ±SEM, Kd values±error estimations are indicated. **(G) –(H)** Interaction of unlabeled titrated human recombinant MANF (0-9.3µM) with Alexa647-labeled through His-tag luminal domain of IRE1α (20nM) in presence of increasing concentrations of Ca^2+^ concentration (100µM-2.5mM). Microscale thermophoresis binding curves, showing mean fraction bond values from n=3 individual repeats per binding pair ±SEM, Kd values±error estimations are indicated.

**Supplementary Figure 2.**
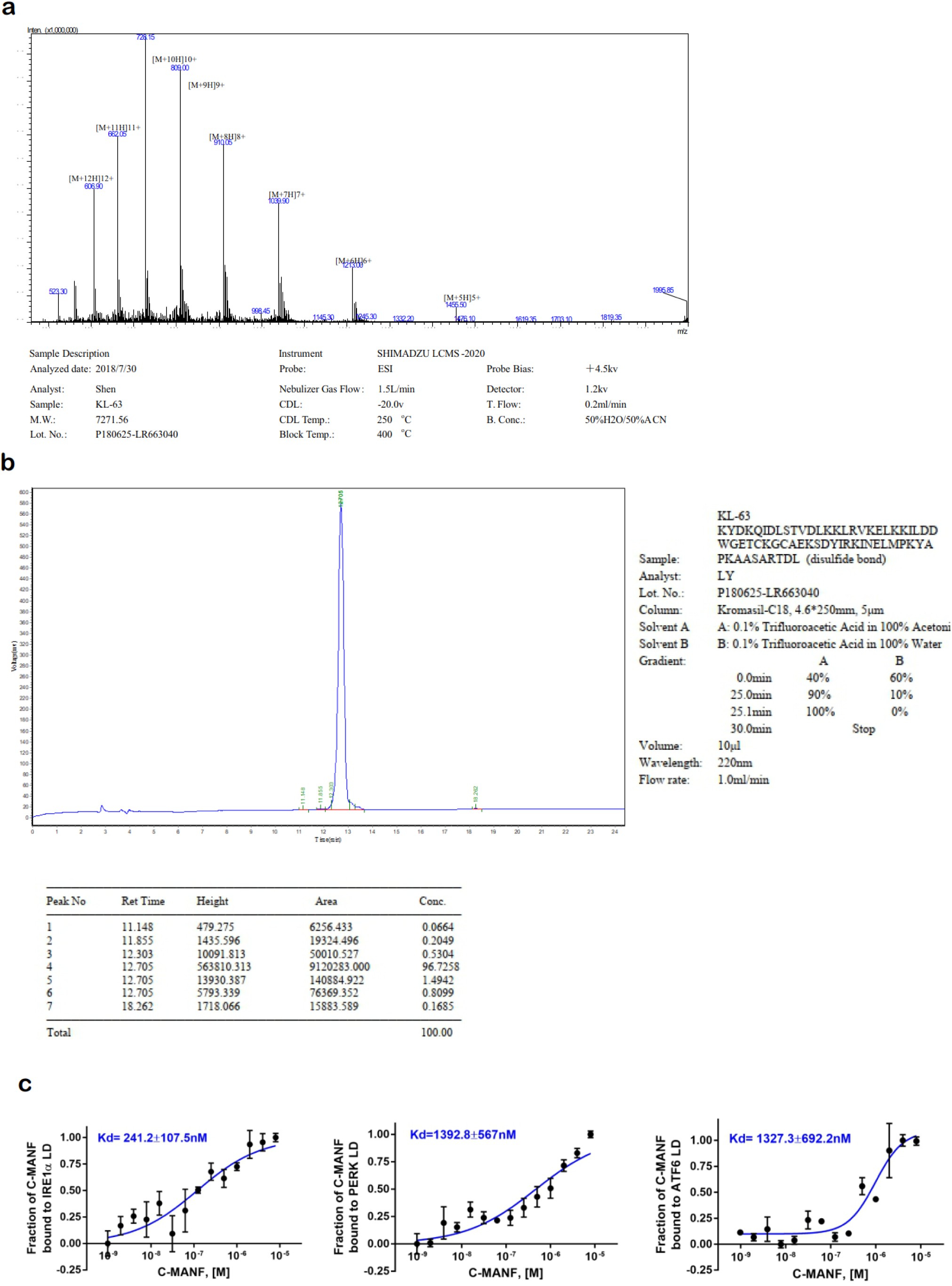
Properties of chemically synthesized and E. coli expressed C-MANF. **(A)** C-MANF forms a single disulfide bond between Cys128 and Cys130 as shown by mass spectrometry (MS) **(B)** C-MANF is homogeneous as shown by high-performance liquid chromatography (HPLC) **(C)** *E*.*coli* produced C-MANF (0-8 µM) is interacting with labeled through His-tag LDs of UPR sensors IRE1α, PERK, ATF6 (20nM). Microscale thermophoresis binding curves, showing mean fraction bond values from n=3-4 experiments per binding pair ±SEM, Kd values±error estimations are indicated.

**Supplementary Figure 3.**
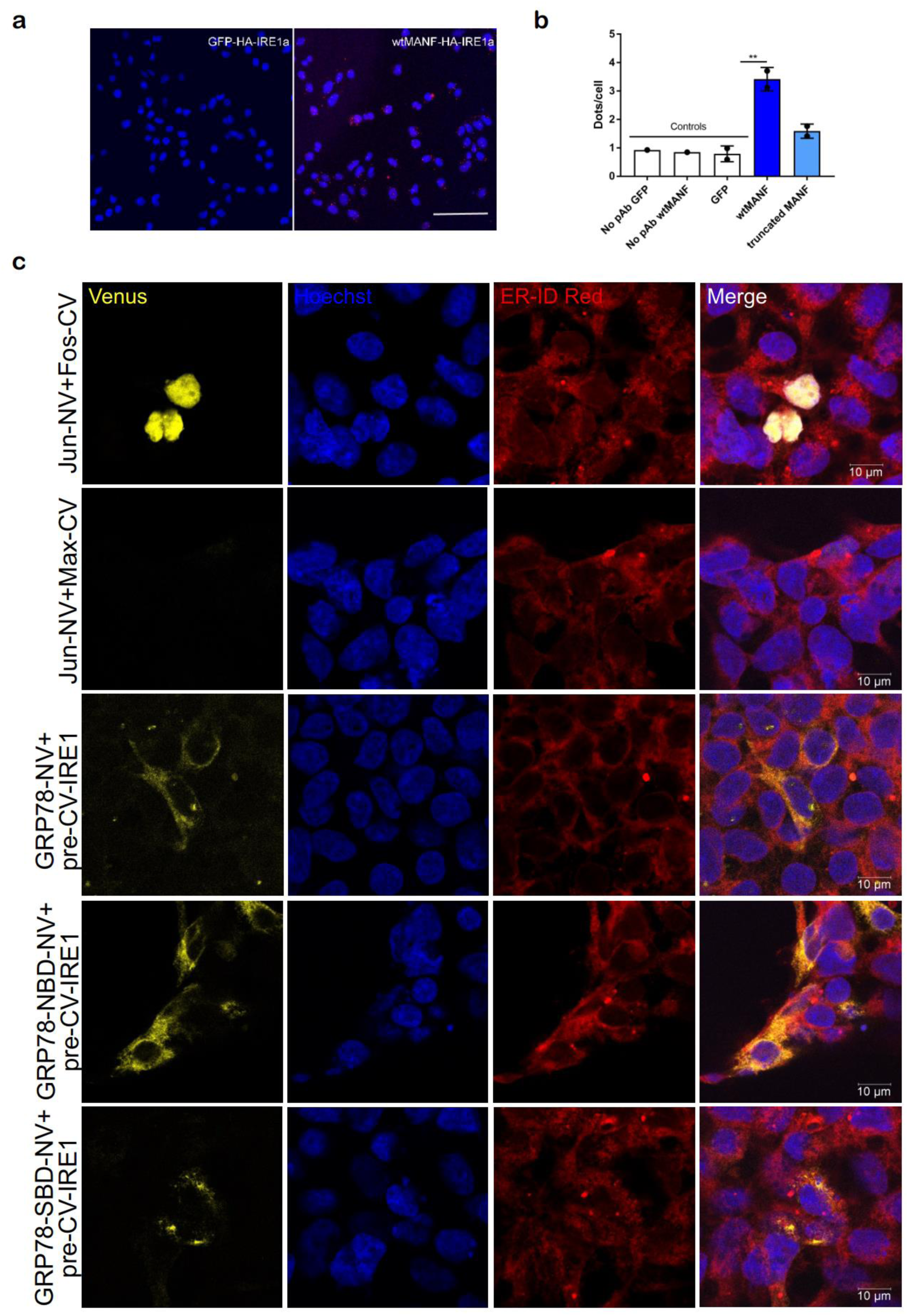
MANF is interacting with IRE1α in CHO cells. **(D)** MANF-HA interaction with IRE1α in CHO cells, shown using Duolink™ proximity ligation assay **(E)** Quantification of MANF interaction with IRE1α in CHO cells, n=3 **(F)** IRE1α is mainly interacting with nucleotide binding domain of BiP (BiP-NBD) and not substrate binding domain of BiP (BiP-SBD). BIFC data, n=3.

**Supplementary Figure 4.**
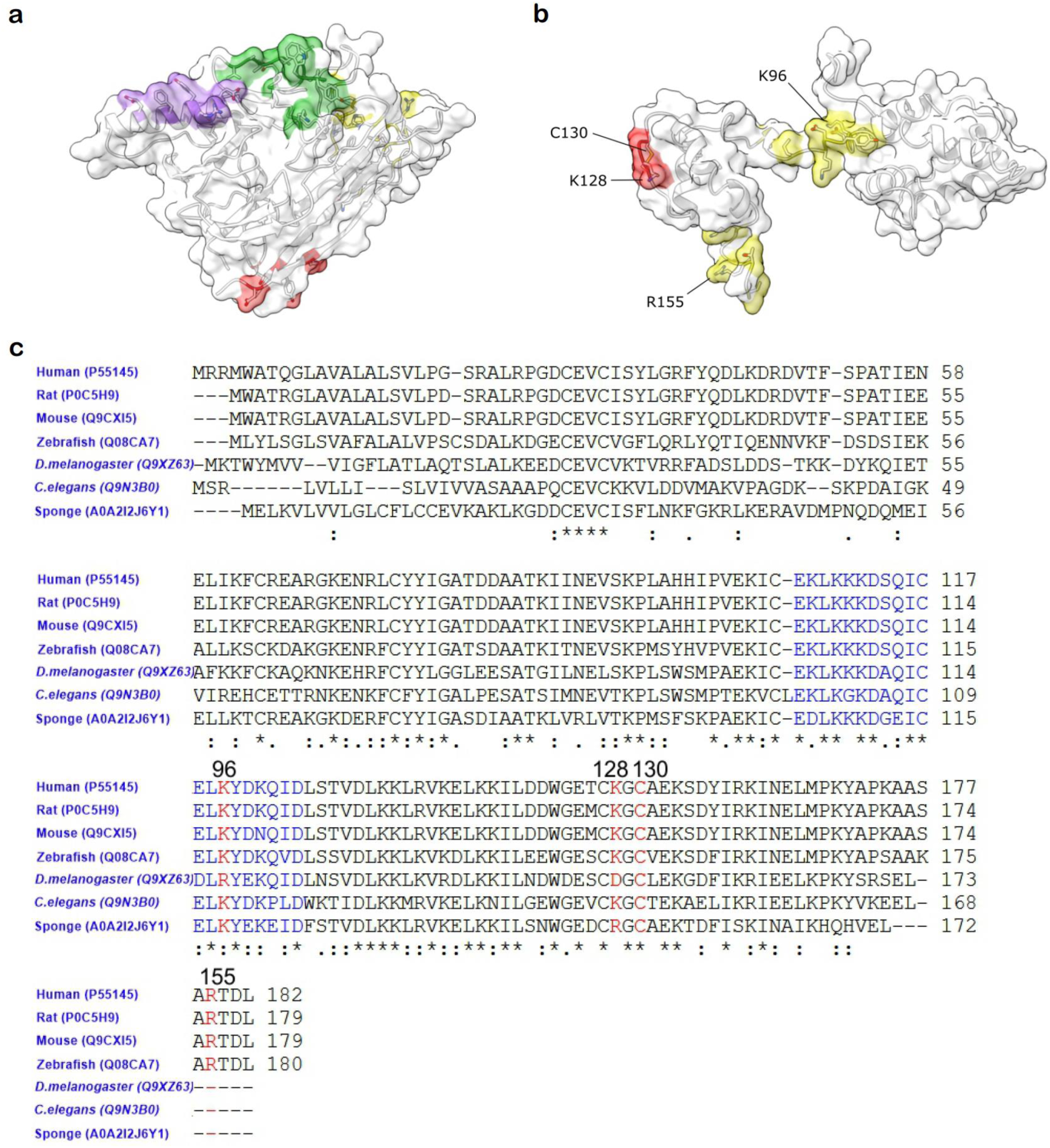
Putative IRE1α binding sites in MANF according to computational modeling. **(A)** Potential binding sites of IRE1α. Labelled amino acid residues of each potential binding site are colored differently. **(B)** Potential binding sites of MANF. Labelled amino acid residues of each potential binding site are colored differently. **(C)** Evolutional conservation of putative IRE1α binding sites in MANF. Mutated amino acid are indicated with red (fuxia) color. Highly conservative region between N- and C-terminus of MANF indicated with blue color.

**Supplementary Figure 5.**
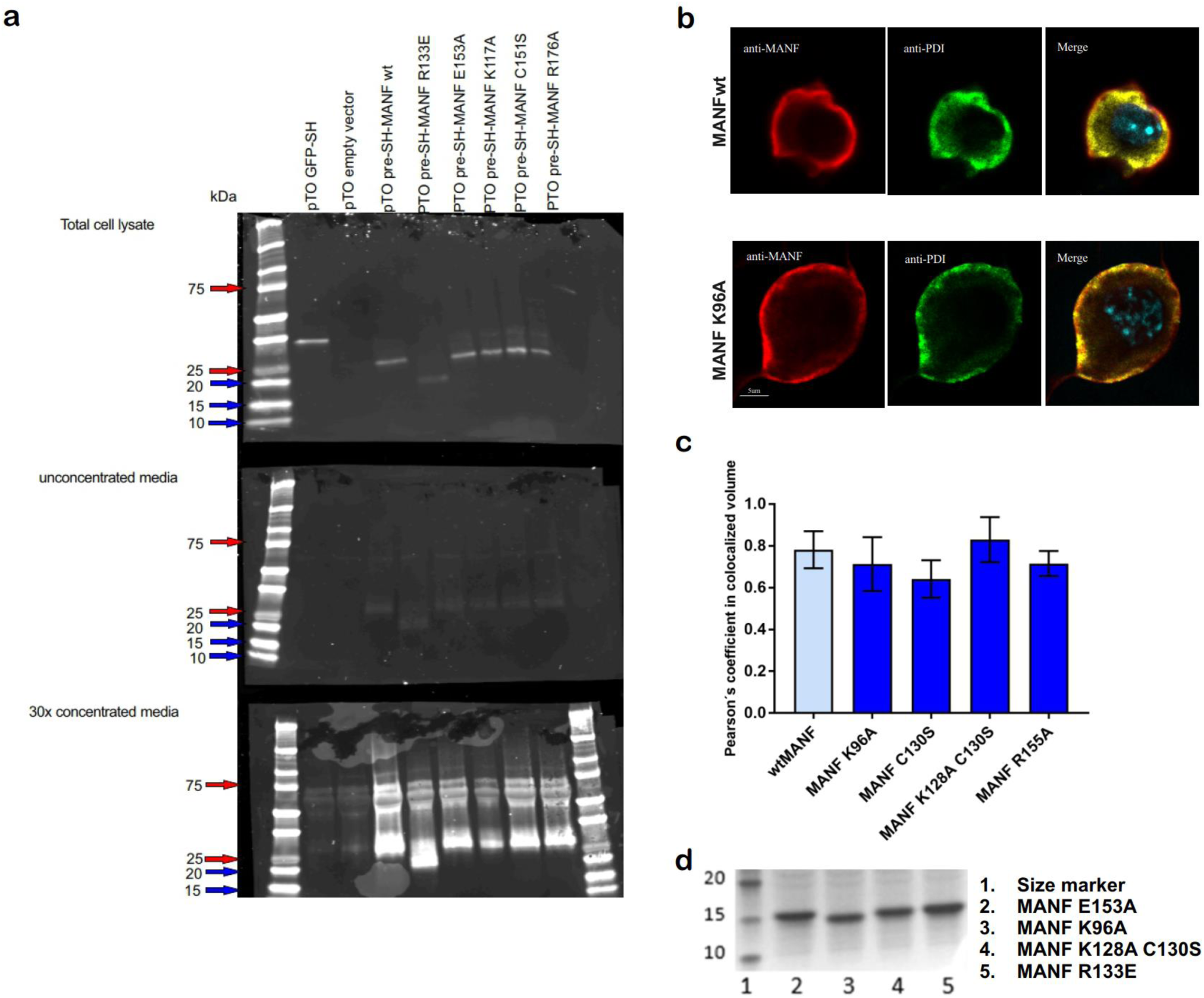
Mutant human recombinant MANF proteins are expressed, secreted and localized in cells similarly to wtMANF. **(A)** Expression and secretion of putatively deficient for IRE1α LD binding MANF mutant constructs in HEK293 cells **(B)** Localization of putatively deficient for IRE1α LD binding MANF mutant constructs in SCG neurons is similar to that of wtMANF construct (ICC). Representative image of wtMANF and MANF K96A, scale bar 5µm. **(C)** Localization of putatively deficient for IRE1α LD binding MANF mutant constructs in SCG neurons is similar to that of wtMANF construct (ICC). Quantification of Pearson’s coefficient in colocalized volume for different MANF mutants, n=5 independent experiments. **(D)** SDS-PAGE gel electrophoresis of purified from *CHO* cells human recombinant MANF K96A and MANF K128AC130S proteins (bands 3 and 4, correspondently), non-reducing conditions, Coomassie blue staining.

**Supplementary Figure 6.**
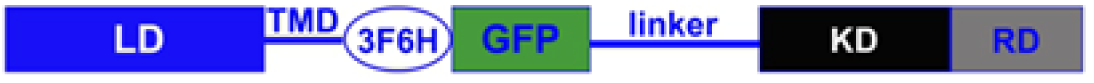
The scheme of IRE1α-3FGH-GFP construct

